# *In situ* Alphavirus Assembly and Budding Mechanism Revealed by Cellular CryoET

**DOI:** 10.1101/2021.10.14.464449

**Authors:** David Chmielewski, Michael F. Schmid, Graham Simmons, Jing Jin, Wah Chiu

## Abstract

Chikungunya virus (CHIKV) is an alphavirus and the etiological agent for debilitating arthritogenic disease in humans. Previous studies with purified virions or budding mutants have not resolved the structural mechanism of alphavirus assembly *in situ*. Here we used cryogenic electron tomography (cryoET) imaging of CHIKV-infected human cells and subvolume classification to resolve distinct assembly intermediate conformations. These structures revealed that particle formation is driven by the spike envelope layer. Additionally, we showed that asymmetric immature nucleocapsids (NCs) provide scaffolds to trigger assembly of the icosahedral spike lattice, which progressively transforms immature NCs into icosahedral cores during virus budding. Further, cryoET of the infected cells treated with neutralizing antibodies (NAbs) showed that NAb-induced blockage of CHIKV assembly was achieved by preventing spike-spike lateral interactions that are required to bend the plasma membrane around NC cores. These findings provide molecular mechanisms for designing antivirals targeting spike-driven assembly/budding of viruses.

## Introduction

Chikungunya virus (CHIKV) is the most common alphavirus infecting humans worldwide, causing epidemics in all continents except Oceania and Antarctica. Transmitted primarily by *Aedes* mosquitoes, CHIKV infection is associated with severe symptoms of debilitating and often chronic polyarthritis in infected individuals (Silva and Dermody, 2017). No licensed vaccine or antivirals are available for CHIKV treatment. The CHIKV virion is ∼70nm in diameter, with a membrane-embedded envelope glycoprotein (GP) shell of 240 copies of E1-E2·E3 heterodimers arranged as 80 trimeric spikes, and an inner nucleocapsid (NC) core of 240 capsid proteins (Cps) that encapsulate the 11.5kb plus-sense (+) genomic RNA (gRNA) (Sun et al., 2013). In CHIKV-infected cells, viral structural proteins (Cp, E1, E2, E3) are synthesized as a single polyprotein precursor molecule. Cp auto-proteolytically cleaves itself from the polyprotein and oligomerizes into nucleocapsid-like-particles (NLPs) in the cytosol, mediated by Cp interactions with the negative-charged gRNA and Cp-Cp interactions (Choi et al., 1991, 1997; Nicola et al., 1999). The remaining polyprotein is inserted into the ER, processed into E1/E2·E3 heterodimers and trafficked through the secretory pathway to the plasma membrane (PM) as trimeric spikes. E3 is cleaved off by host furin and furin-like proteases and stays associated with the nascent virions (Basore et al., 2019; Zhang et al., 2011). Convergence of cytosolic NLPs and membrane-embedded spikes at the cell surface results in assembly and budding of enveloped virions with icosahedral spike and NC layers (Cheng et al., 1995).

Formation of enveloped virus particles requires viral protein and/or host factor-induced curving of a cellular membrane around viral cores followed by membrane scission. Some viruses such as retroviruses, rhabdovirus and filoviruses, recruit host ESCRT machinery to drive virus assembly/budding and their NC cores alone can bud as virus-like particles (VLPs) without viral glycoproteins embedded in the lipid envelope. In contrast, production of CHIKV VLPs requires co-expression of GPs and Cp, while budding is reported to be ESCRT-independent (Noranate et al., 2014; Taylor et al., 2007). Numerous studies suggest that alphavirus budding is mediated by both vertical spike:Cp interactions and lateral interactions between spikes (Forsell et al., 2000; Suomalainen et al., 1992). Insertion of the intracellular tail of E2 into the hydrophobic pocket of Cp C-terminal domain mediates the vertical Spike-Cp interactions, while lateral E1 self-interactions form the surface envelope lattice (Cheng et al., 1995; Tang et al., 2011; Zhang et al., 2011).

Two distinct models for the major driving force in budding have been proposed: spike-NC binding and spike-spike shell interactions (Garoff et al., 2004). Purified Cps assemble *in vitro* into core-like-particles (CLPs) with fragile, imperfect icosahedral symmetry (Mukhopadhyay et al., 2002; Wang et al., 2015). Microinjection of *in vitro*-assembled CLPs into spike-expressing cells induces low levels of virus budding, suggesting interactions between spikes and preassembled cytosolic NLPs drive budding (Cheng and Mukhopadhyay, 2011; Snyder et al., 2011). In support of the alternative spike-driven model, Cp mutants deficient in Cp-Cp interactions can still form virions with viral spikes, though at a lower efficiency than the wild-type (wt) (Forsell et al., 1996). This suggests that spike-spike interactions and spike-Cp interactions are sufficient to drive the assembly and budding of icosahedral particles without a preassembled NC. There are also reports of propagation of capsidless alphavirus via infectious microvesicles at a titer logs lower than the wt virus (Ruiz-Guillen et al., 2016). How spikes mediate packaging of gRNA in membrane vesicles is largely unknown. It is unclear whether any of these budding models, derived from viral mutants, are applicable to the case of wt alphavirus budding *in situ*.

Our previous study demonstrated that neutralizing antibodies (NAbs) inhibit CHIKV budding by crosslinking spikes at the outer surface of the PM and trapping NLPs inside CHIKV-infected cells (Jin et al., 2015, 2018). It suggests that spike organization on the infected cell surface is critical for virion assembly and budding. What is left unaddressed in that study are the molecular details of NAb-bound spike organization that inhibit membrane bending. Defining the difference between the spike organization in the normal virus budding process and that of budding inhibitory spike-NAb complexes can inspire future therapeutic antiviral strategies. In the same study, we reported lack of icosahedral symmetry in budding-arrested NLPs and suggested it could either be a structural feature of immature cytosolic NLPs or a result of disassembly of budding-arrested NLPs. NC assembly is a potential antiviral target and necessitates further characterization of NC morphogenesis prior-to and during budding (Wan et al., 2020). Recently, Cp-gRNA interactions were found to differ between cytosolic NLPs and virion-NCs (Brown et al., 2020), supporting previous reports of physical differences between the two populations and suggesting currently-uncharacterized morphological changes occur during virus budding (Lamb et al., 2010).

To determine the assembly events leading to CHIKV mature particle formation, we utilized cryogenic electron tomography (cryo-ET) imaging and subtomogram averaging to reveal structures of intermediately assembled particles at multiple stages of virus budding in CHIKV-infected human cells. Additionally, we resolved structures of self-assembled spikes and NLPs in the absence of spike-Cp interactions. Based on these structures, our study defines the mechanistic roles of spikes and immature NCs in CHIKV budding and elucidates conformational changes of NC during virion assembly. In addition, we present a method of classifying snapshots of the alphavirus budding process into discrete ensemble averages of conformational states *in cellulo*.

## Results

### CryoET and subtomogram classification of CHIKV budding intermediates *in situ*

CHIKV particle assembly and budding, driven by interactions between NCs and membrane-embedded spikes, is known to occur at the PM of infected cells. To capture the dynamic process in the near-native state, we imaged U2OS cells, a human bone osteosarcoma cell line, that were infected with CHIKV-181 vaccine strain and embedded in vitreous ice. Collection of tomographic tilt series at the infected cell peripheries 8 hours post-infection revealed a variety of budding phenotypes (movie S1), with CHIKV assembly events located at the PM of the cell body (Fig. 1A-D), on long intercellular extensions (>10 µm), short extensions (typically 2-10 µm in length), and thin branching extensions composed solely of viral particles (Fig. 1A-B). Particles at various intermediate stages of budding at the PM, as well as fully assembled virions released into the extracellular space, were observed (Fig. 1E-F), thus capturing snapshots of the entire CHIKV assembly/budding process. The high concentration of CHIKV budding at specific regions of the cell periphery and variety of observed cell extensions is consistent with previous reports of highly localized budding and virus-induced branching structures at the cell periphery from 2-D electron microscopy images of plastic-embedded material (Birdwell et al., 1973; Laakkonen et al., 1998; Martinez et al., 2014; Pavan et al., 1987). Interestingly, CHIKV replication spherules, where viral RNAs are synthesized, were occasionally observed near cytosolic NLPs and budding viruses (Fig. 1C-D, Fig. S1). It is conceivable that viral RNAs are synthesized and immediately packaged into NLPs that bud into virions, all near the PM.

**Figure 1.**
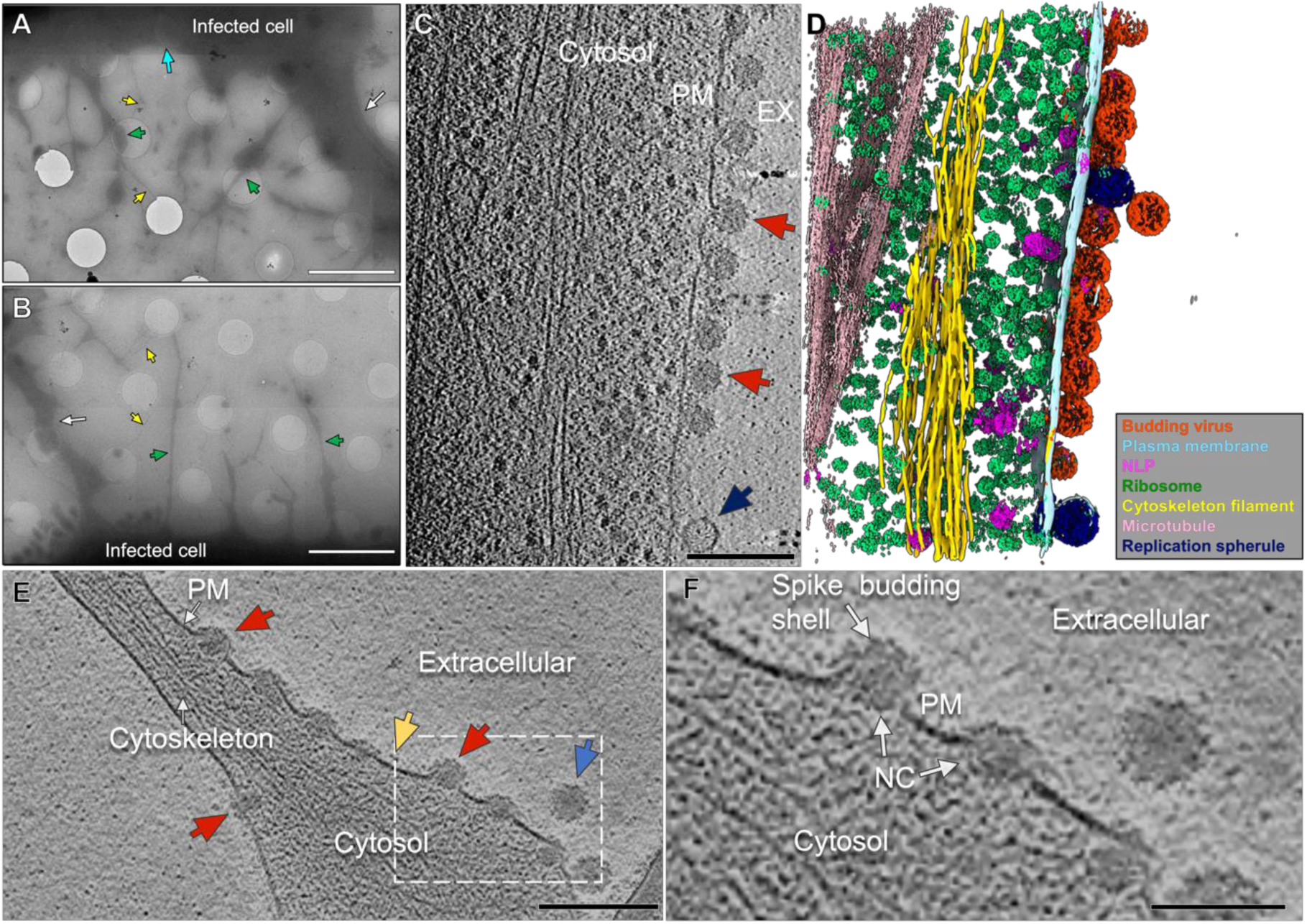
CHIKV assembly and budding at the infected-cell periphery. (A-B) Low magnification images of the cell periphery reveal cell body (cyan arrow), long intercellular extension (white arrow), short extensions enriched in virus assembly features (green arrows), and thin extensions of viral particles (yellow arrows) emanating from the short extensions or cell body. Scale bars 5 µm. (C) Tomographic slice of cell periphery depicting virus budding events (red arrows) and RNA replication spherules (navy blue arrow) at the PM with (D) corresponding 3D segmentation of cellular features. (EX: extracellular space) Scale bar 200 nm. (see also Fig. S1) (E) Tomogram slice of short extension with budding intermediate particles (red arrow), spikes (yellow arrow), and cell free virion (blue arrow). Scale bar 200 nm. (F) Enlarged view of the boxed region in (E) shows intermediate viral assembly complexes at the PM, composed of a spike budding shell and nucleocapsid (NC). Scale bar 100 nm.

On the virus-infected cell periphery we frequently identified thin extensions formed by incomplete particles, often the width of just a single virion (∼70 nm diameter, <5 µm length) and lacking bundled cytoskeleton filaments (Fig. S2). Thin extensions possess a diversity of particle structures, with differences in the levels of completion and structural conformations (Fig. S2). Convergence of two opposing membranes containing near-continuous budding intermediates was observed to give rise to strings of incomplete particles with a continuous membrane connection. Due to the lack of sufficient spikes to finish enclosing the NC as an icosahedron, these linked particles are unlikely to complete the assembly of full virions and therefore were excluded from the following analysis.

In order to determine *in situ* conformations of the CHIKV particle assembly process, we extracted 1,918 budding intermediate particles at the cell periphery (Fig. 2C, 3A). To address conformational and compositional heterogeneity within snapshots of the budding process, individual particle subvolumes were discretized into 3D classes through an unbiased and iterative multi-reference classification based on structural similarity using C5 symmetry (Fig. S3A, see “Methods” section). This resulted in 12 3D reconstructions of discrete virus budding conformations (Fig. 2) that displayed progressive levels of budding, with different extents of completion of the spherical budding shell. The 3D class averages 4, 6, 9 and 10 possessed weak density of the PM and trailing end of NC that became blurred and masked-out during refinement. This is likely due to heterogeneity among individual particles within those classes and/or lack of symmetry of these regions relative to the leading end of budding particles.

**Figure 2.**
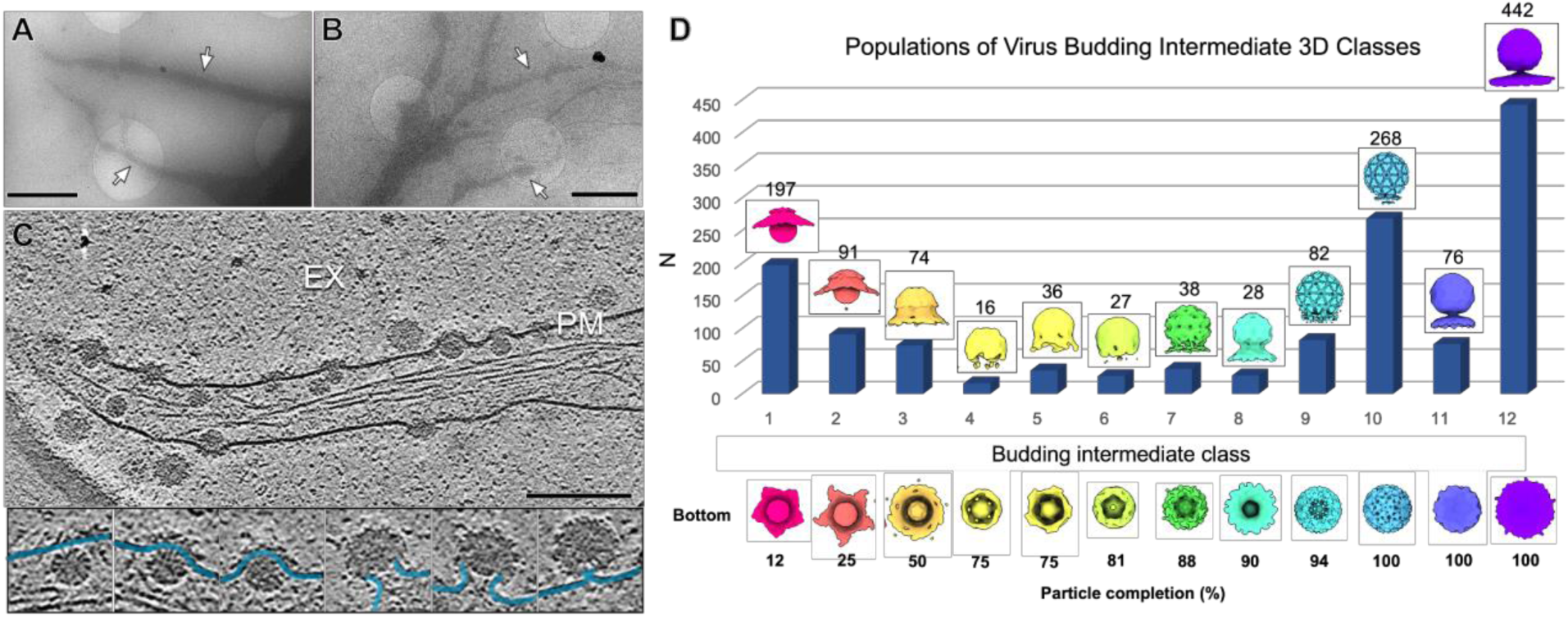
Classification and enumeration of CHIKV budding intermediates. (A-B) Images of virus-infected cells with extensions enriched in virus assembly (white arrows). Scale bars 2 µm. (see also Fig. S2) (C) Tomogram slice image depicts snapshots of the virus budding process. Selected particle images (below) reveal heterogeneity based on conformations of the bending PM (blue). (EX-extracellular space), Scale bar 200 nm. (D) CHIKV-budding intermediate 3D classes determined by subvolume classification. Density maps of each class average (1-12) are colored uniquely and displayed with side-view and bottom-view (viewed from below the PM). The number of particles (N) assigned to each class displayed as a bar graph with respective N listed above. (see also Fig. S3)

**Figure 3.**
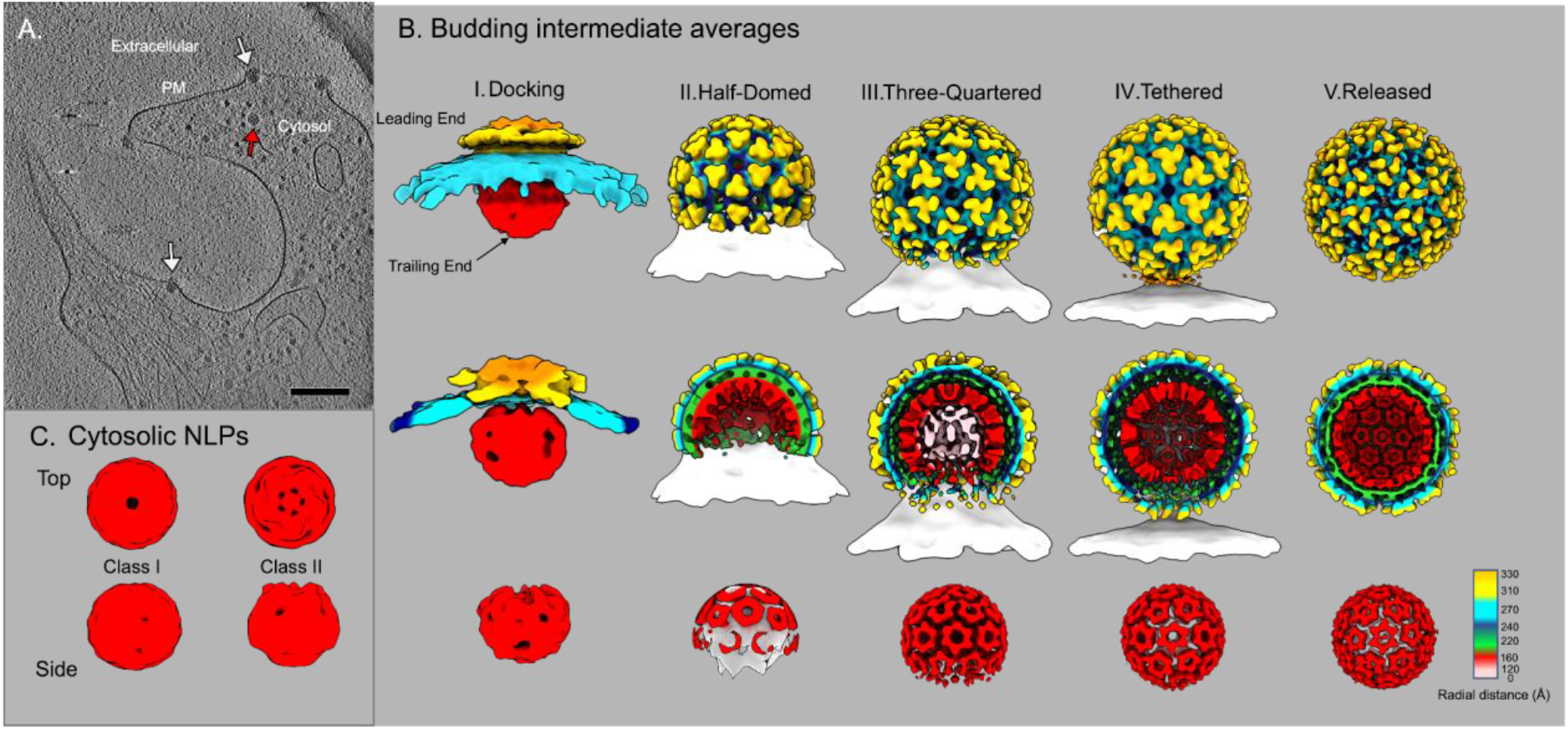
Refined structures of CHIKV assembly intermediates. (A) Tomogram slice image displays budding intermediate particles at the PM (white arrows) and an apparently cytosolic NLP (red arrow). Scale bar 200 nm. (B) Radially-colored density maps of five CHIKV ensemble subtomogram averages arranged in accordance with budding progression from earliest budding level (“I.Docking”) to latest (“V.Released” virion) with half-cut representations (middle row) and NC-zoned densities (bottom row). PM and NC density (“Half-Domed”, bottom row) from class averages prior to additional 3D refinement depicted as white surfaces. (see also Fig. S3 & S4) (C) Subtomogram average structures of two cytosolic NLP classes. Class I displays no interpretable 5-fold symmetry while class II shows weak five-fold symmetry at one pole. (also see Fig. S4 & S5).

To analyze progression of the CHIKV budding pathway, the population of particles within each 3D class was then analyzed in relation to budding level (i.e., percentage completion of the budding shell). Budding level was determined by using the mature icosahedral virion as a reference and counting the number of spikes (out of 80) covering the budding shell of each low-resolution intermediate class average (Fig. 2D). From a total of 1,375 budding intermediate particles within the 12 classes that converged to interpretable 3D structures, 288 particles (21%) were classified into structures containing 10-20 spikes (classes 1-3,12-25% complete), 191 particles (14%) were classified into structures containing 40-70 spikes (classes 4-8, 50-88% complete), and 868 particles (63%) were classified into structures containing 75-80 spikes (classes 9-12, 94-100% complete). The low particle numbers and comparatively low resolution classes in the 50-88% of completed virion range (budding classes 4-8) suggest CHIKV assembly progression at these mid stages is more transient than early (budding classes 1-3) and late stages (budding classes 9-12).

To enhance low-resolution image contrast for visualization of CHIKV particles, we performed another set of experiments which utilized a Volta phase plate (Danev et al., 2014). Previous structural studies of purified alphavirus particles have described significant inter- and intra-particle heterogeneity (Chen et al., 2018; Zhang et al., 2011) but concerns about the fragility of enveloped viruses to purification have cautioned conclusions about the relevance of structural heterogeneity to alphavirus assembly *in situ*. Therefore, our imaging of CHIKV budding and released virions *in situ* eliminates the need for *in vitro* purification and handling of virus particles prior to vitrification. Within released virions, relatively absent density was consistently observed at one side of the particle between spike and NC core layers (Fig. S4). Further, unidentified molecular complexes are observed at the base of the V-shaped viral envelope in the relatively absent density region of many released particles (Fig. S4D). Relatively absent density was also observed in released multi-core particles in the region between NCs, suggesting it arises from a lack of spike-NC contacts enclosing the icosahedron (Fig. S4D-E). Interestingly, the trailing-end of late-stage budding particles tethered to the PM displays similar relatively absent density and geometry of the viral envelope (Fig. S4B). Therefore, deviations from the icosahedral lattice within released virus particles appear to directly result from late events in virus budding from the host cell membrane.

### *In situ* structures of progressing CHIKV budding intermediates

To further analyze structures of spike:immature NC complexes at the molecular level during particle assembly, we performed additional 3D refinement of released virions and four budding-intermediate class averages that displayed low-resolution icosahedral features (see Methods). Icosahedral (5-3-2) symmetry is a feature of purified CHIKV particles (Sun et al., 2013), but it was unknown if CHIKV assembly progressed through partially-icosahedral intermediates or if full icosahedral symmetry of infectious particles arose from protein rearrangements following virus release. Particle subvolumes were aligned with C5 symmetry, resulting in the earliest budding structure displaying a 5-fold pentagon at the leading end of budding, while the other four structures displayed excellent pentagon and hexagon assemblies with 5-fold, 3-fold, and 2-fold symmetry axes (Fig. 3B). The five subvolume averages of discrete budding states range in resolution from 8.3 Å (released) to ∼44 Å (“docking”) (0.143 FSC criterion) (Fig. 3B, Fig. S3B). Subnanometer resolution in the released virion average is validated by the visualization of transmembrane helices of E1/E2 (Fig. S3C-D). The differences in resolution among the budding intermediate ensemble averages are likely caused by increased conformational flexibility in those assembly states with less icosahedral symmetry constraints and compositional differences between aligned particles with slightly different budding levels.

Within the budding intermediate averages, a striking correlation between icosahedrally-symmetric regions of the spike budding shells and their underlying NC core was observed at the “leading end” of budding (Fig. 3B). In contrast, there is a lack of detectable symmetry in the “trailing end” of each intermediate NC’s structure, where no spikes are present (Fig. 3B, Fig. S4A-C). Even within the latest-stage budding conformation (“tethered”), weak density of the final penton of spikes at the trailing end is matched by a disordered pentamer of Cps in the NC below it (Fig. S4C). Based on these structures, as the icosahedral spike shell grows during budding, it appears to reorganize those Cps of the immature NC below into matching icosahedral symmetry through 1:1 spike:Cp interactions. This result revealed that assembly progresses through partially icosahedral intermediates and explains the origin of matching T=4 icosahedral spike and NC lattices in released alphavirus virions (Cheng et al., 1995).

In the NC-centric model of alphavirus budding (Garoff et al., 2004), preformed icosahedral NLPs would provide a symmetric template for incorporation of spikes at the cell surface. In our study, 545 subvolumes of cytosolic NLPs without clear attachment to spikes at the PM were analyzed for any icosahedral symmetry that could direct virion assembly and budding (Fig. 3A, Fig. S5A). Following 3D-classification, respective sets of NLPs were refined with C5 symmetry, resulting in two NLP averaged structures (class I & II), both lacking icosahedral symmetry (as seen in released virion NCs) or clear 5-fold symmetry of Cps (identified at the leading ends of budding intermediate NCs) (Fig. 3C, Fig. S4A). However, cytosolic NLP class II does display weak five-fold symmetry at one pole, raising the possibility that those NLPs were already interacting with spikes at the PM but their orientations in the tomograms prevented that observation. In addition, both cytosolic-NLP structures and the NC of the earliest “docking” budding-intermediate are significantly smaller than spherical NCs of the latest-stage budding intermediate (“tethered”) and released virions (diameters ∼43 nm) (Fig. S4A). The NC of the “docking” budding intermediate and cytosolic NLP exhibiting weak five-fold symmetry at one end (class II) are both oblate spheroids, with long axis of ∼37 nm and short axis between 31 nm and 33 nm, respectively (Fig. 3B-C, Fig. S4A).

The cytosolic NLP structure with no observable five-fold symmetry (class I) is roughly spherical with diameter ∼37 nm (Fig. 3C, S4A). For class I cytosolic NLPs, our result does not rule out that Cps can be arranged in assemblies with other non-icosahedral or non-five-fold symmetry. It is also possible that the low contrast of NCs in the cytosol results in inaccurate alignments and a non-symmetric average, though we consider this unlikely because of the following observations. To determine if weak five-fold symmetry in the cytosolic NLP class II average was located randomly on individual particles or the result of Cp:spike interactions, we mapped the refined subvolume particle orientations in 3D back to the originating tomograms. This confirmed that most particles within class II were positioned such that the five-fold organized-pole of the NLP was oriented towards the PM, presumably interacting with membrane-embedded spikes (Fig. S5). Weak five-fold symmetry within NCs prior to budding likely results from Cp:spike contacts at the PM, while the rest of the NLP structure lacks five-fold or icosahedral symmetry. Therefore, true cytosolic NLPs (class I) are structurally heterogeneous and lack icosahedral or local five-fold Cp organization. During budding, immature NLPs must undergo a significant structural maturation from an initial structurally heterogeneous RNA-Cp assembly to expanded, near-icosahedral viral NCs following ordering interactions with the icosahedral spike lattice.

### Assembly of spike lattices

Structures of the budding-intermediates revealed a progressive spike-driven NC morphogenesis, demonstrating the importance of the spike lattice in CHIKV particle assembly. However, it was not known how the spike lattice acquires icosahedral symmetry that reorganizes the NC. One possibility is that individual spikes bind to NCs during budding and form lateral interactions with other spikes on the budding particle surface (Forsell et al., 2000). Another proposal of alphavirus budding from preformed, higher-order spike assemblies on the PM arose from previous reports of hexagonal spike lattices as two-dimensional planes or 6-fold helical tubes (von Bonsdorff and Harrison, 1978; Kononchik et al., 2009; Soonsawad et al., 2010). These alternative spike assemblies were generated either by treating virions with detergent (von Bonsdorff and Harrison, 1978) or mutating both E1/E2 GPs (Kononchik et al., 2009), or observed inside cytopathic vacuoles in virus-infected cells (Soonsawad et al., 2010). Whether or not the alternative spike assemblies form at the PM of wt alphavirus-infected cells, and their relevance to wt alphavirus assembly/budding, was unknown. In our study, we visualized both spike organization near budding intermediates at the PM (Fig. 4), and structures of self-assembled spikes in wt CHIKV-infected cells *in situ* (Figs. 4&5).

**Figure 4.**
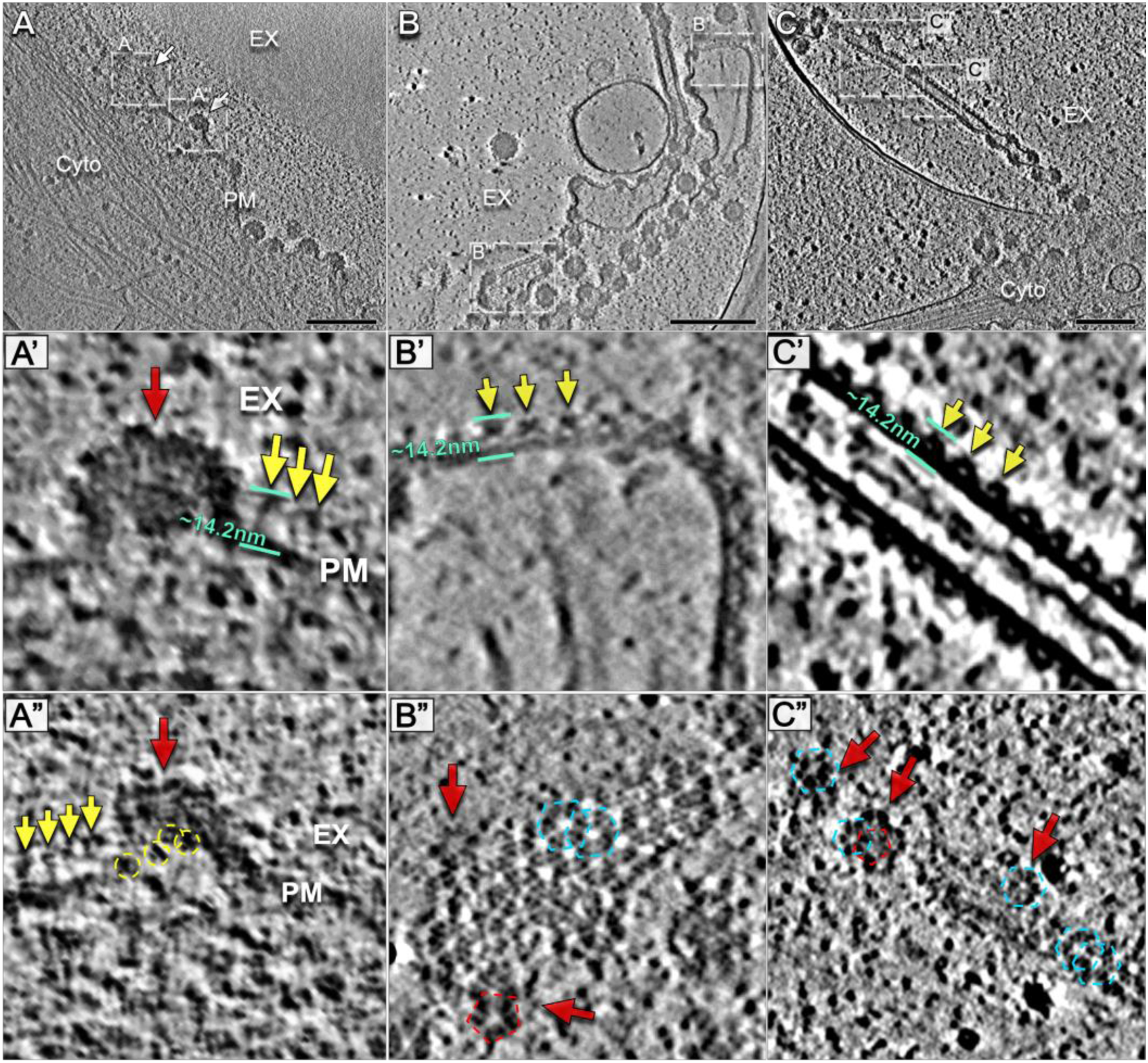
Spike organization on the virus-infected cell surface. Volta phase plate tomogram slice images of (A) cell periphery with budding intermediate particles (white arrows) and (B,C) thin cell extensions enriched in linked, early-stage incomplete particles. Thin cell extensions include regions of planar sheets of spikes (B, dashed white boxes) and tubular array of spikes (C, dashed white box). (A’) Detailed tomogram slice view displays apparent spikes (yellow arrows) near a budding particle (red arrow). The distance between the crest of apparent spikes and the inner leaflet of the PM (aquamarine lines) is measured to be ∼14.2 nm. (A’’) Slice image displays apparent spikes as side-views (yellow arrows) and top-views (yellow circles) near base of budding particle (red arrow). (see also Fig. S6) (B’,C’) Detailed tomogram slice displays side views of PM with laterally organized spikes (yellow arrows) in planar sheets and helical tubes, respectively. Distances between spike crest and inner leaflet of PM again displayed in aquamarine. (B’’) Detailed tomogram slice with top view of a planar sheet reveals spikes organized as a hexagonal lattice (blue dashed hexagons), with disruption in lattice near budding virus particles (red arrows). Pentagon of spikes identified above NC (red dashed pentagon). (C’’) Detailed tomogram slice of a helical spike tube-like structure reveals hexagonal array of spikes (blue dashed hexagons), while both pentagons (red dashed pentagon) and hexagons are observed on nearby linked, incomplete particles (red arrows).

**Figure 5.**
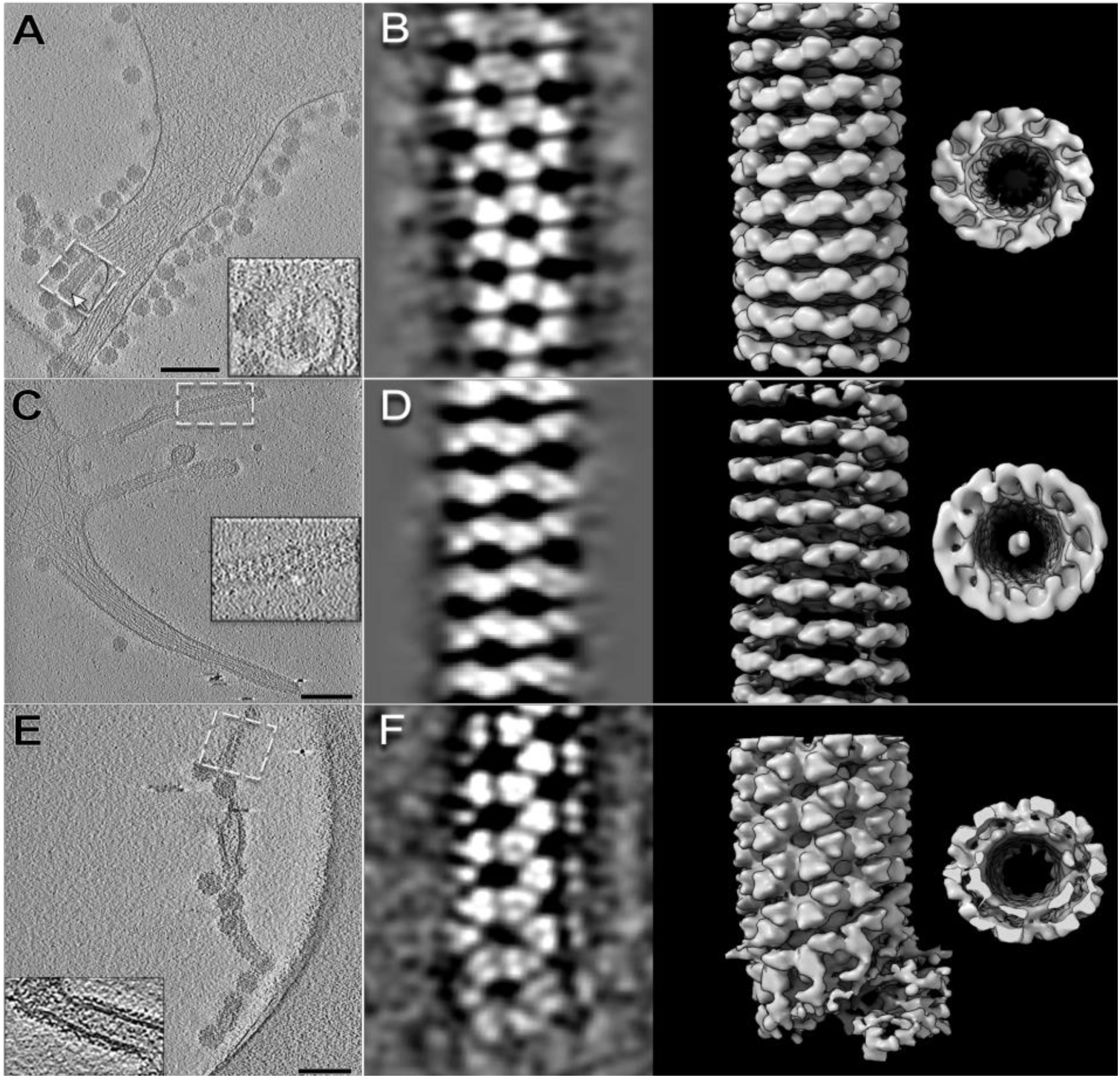
CHIKV envelope spikes arranged in hexagonal lattices form helical tubes in situ. (A,C,E) Tomogram 2D slice images of cell extensions with membrane-embedded spike arrays (white dashed boxes) with enlarged inset views. Arrays form (A) at the base of a budding intermediate particle with NC at the leading end (white arrow), (C) on a cell extension without nearby NCs (see also Fig. S7), and as a (E) segment within a thin extension containing nearby budding viral particles. Scale bars 200nm. (B,D,F) 2D slice view of envelope lattice 3D subtomogram average (left) and density map (middle) (corresponding to A,C,E dashed white boxes) reveal helical arrays of trimeric spikes arranged as hexagon lattices. Density maps of each tube, rotated to view down the helical axis (right), revealed no underlying NCs below the spike lattice and membrane bilayer.

Spike organization at the PM prior to budding has never been observed in virus-infected cells due to the challenges in resolving relatively small macromolecules (∼360kDa) in different orientations in the unstained cell membrane with heterogeneous composition and high image background. In Volta phase plate cryoET data we were able to identify proteins near budding intermediates, potentially individual trimeric spikes based on shape and size, with no discernable high-order organization (dimer-, pentamer-, or hexamer-of-spikes) (Fig. 4A’, A’’). The lack of strict lateral organization between these proteins near the budding shells is significantly different from the spike organization identified in hexagonal spike lattices and the icosahedral-symmetric end of budding particles (Fig. 4B-C, Fig. S6).

Rare instances of near-planar sheets within thin extensions containing high density of spikes (Fig. 4B) were formed by hexagonal spike arrays without underlying NCs (Fig. 4B-B’’). Side-views of spikes in the sheet lattices displayed characteristic spacing and conformation (Fig. 4B’). In addition, highly-curved tubular spike arrays were observed in multiple cellular contexts: at the base of budding particles (Fig. 5A), on extensions entirely devoid of NCs (Fig. 5C, Fig. S7), and on thin extensions loaded with budding particles (Fig. 4C, 5E). Subvolume averages of these tubes, achieved by applying helical rotations to compensate for the tomographic missing wedge, revealed helical organization of trimeric spikes arranged as hexagons (Fig. 5). No internal NC was observed inside any tube. Interestingly, the average diameter of the helical tubes varies from 55 to 65 nm, close to the diameter of the icosahedral CHIKV virion (∼70 nm) (Table S2). Pentagons of trimeric spikes were only observed on the surface of budding-intermediate particles (Fig 4B”, C”), while hexagons were observed on budding particles, helical tubes (Fig. 4C’’, 5) and planar sheet lattices without interior NCs (Fig. 4B’’). The correlation between underlying NC cores and pentagons of trimeric spikes in the envelope lattice suggests that NC cores function to either directly promote five-fold assembly of spikes and/or trigger membrane curvature formation for icosahedral assembly.

Interestingly, compared with the spikes assembled into icosahedral or hexagonal lattices, individual spikes at the base of budding particles not only lack ordered lateral arrangement but also appear highly heterogeneous in shape and orientation (Fig. 4A-A’’, Fig. S6). This suggests that spikes without lateral interactions and spikes assembled into icosahedral or hexagonal lattices have different conformations. These results explain why different epitope residues for specific mAbs were identified using cell-surface displayed spikes, alphavirus-spike pseudotyped HIV-1 reporter viruses, and live alphaviruses (Jin et al., 2015; Kim et al., 2021; Selvarajah et al., 2013). This could suggest that spikes displayed on the surface of spike-transfected cells and pseudoviruses have the same conformation as the individual spikes on the surface of virus-infected cells. Future work leveraging advanced techniques to improve resolution of heterogeneous biomolecules on noisy cellular background is warranted to resolve the conformation(s) of individual spikes on the cell surface before virus assembly.

### Disrupting spike organization to block CHIKV budding

We previously reported that in addition to traditional NAb function in inhibiting virus entry, bivalent binding of NAbs to spikes at the outer surface of CHIKV-infected cells induces coalescence of spikes that inhibits virus assembly/budding (Jin et al., 2015, 2018). To reveal if crosslinking NAbs induce budding arrest by disrupting self-assembly of spikes and/or icosahedral co-assembly of spikes and NC, we imaged CHIKV-infected cells treated with CHIKV-specific NAb C9 (Jin et al., 2015). Large numbers of cytosolic NLPs were observed docking to the inner leaflet of the PM without virus budding (Fig. 6, movie S2), consistent with what we reported previously (Jin et al., 2018). Budding-arrested NLPs are characteristically flattened at the docking end below the near-planar PM, where direct interactions are observed with cytosolic tails of C9-crosslinked spikes (Fig. 6B, G). Side views of the PM revealed a layer of spike ectodomains above the lipid bilayer, and a second layer of dense protein density above the spikes, approximately 150-250 Å from the inner leaflet of the PM (Fig. 6A-B). Proteins above the spike ectodomains, not seen in the regular CHIKV-infected cells, are presumably C9 IgGs bound to the previously reported epitope at the crest of spikes (Jin et al., 2015).

**Figure 6.**
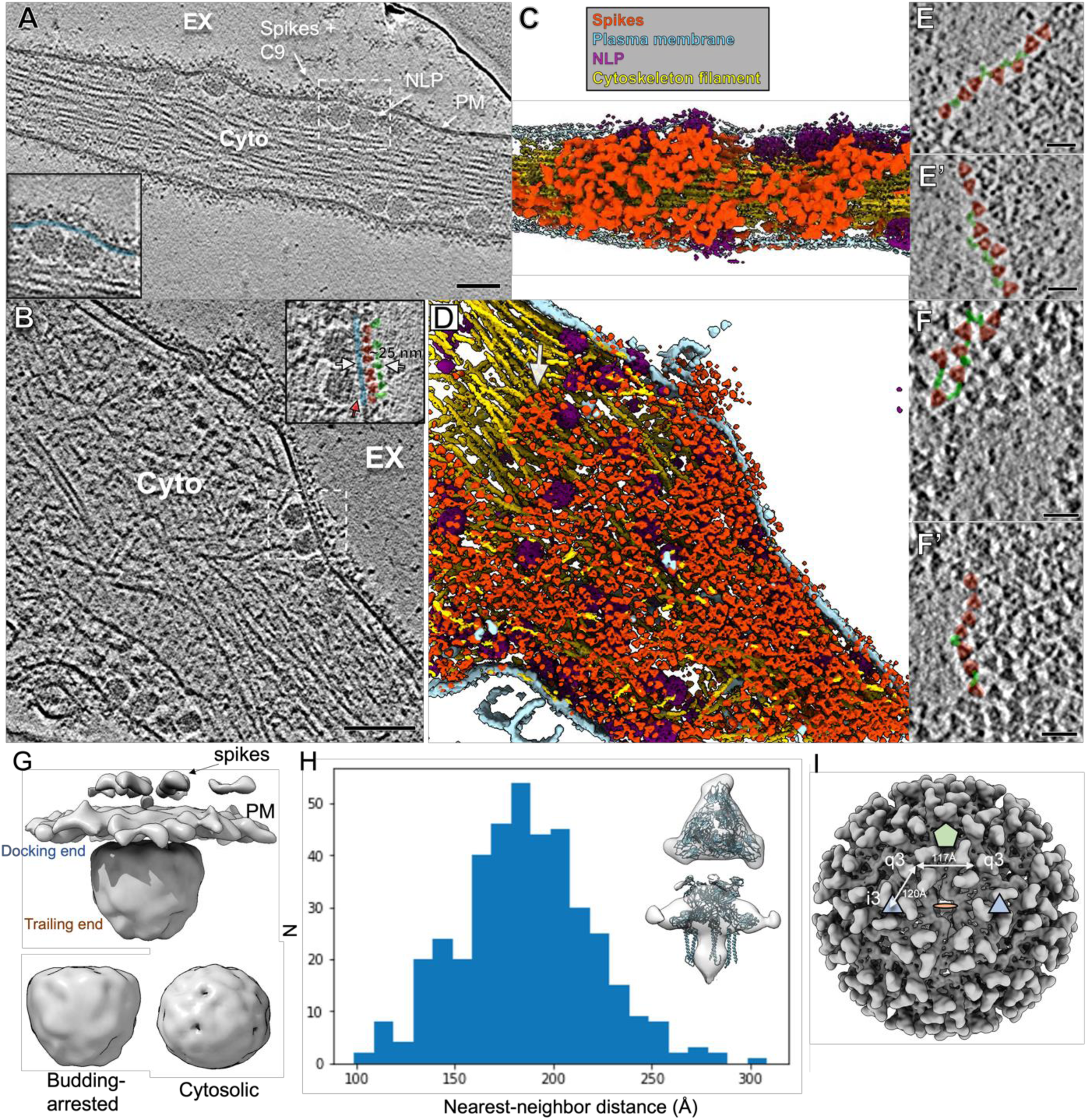
Neutralizing antibody C9 crosslinks spikes at the cell surface and induces coalescence of spike-C9 complexes. (A-B) Individual slices of Volta phase plate cryoET tomograms displaying CHIKV-infected cells treated with NAb C9 revealed arrested NLPs at the PM inner leaflet and dense, coalesced spike-C9 complexes on the PM outer leaflet (inset images: docked NLPs interacting with spike intracellular tails (red arrow) at the PM inner leaflet (blue), with spike ectodomains (pink) bound to NAb C9 (green) outside the cell. Scale bars 100nm. EX-extracellular, Cyto-cytosol. (C-D) Corresponding 3D cellular annotations of tomograms (A-B), with trimeric spikes (orange-red), PM (light blue), NLPs (purple) and cytoskeleton filaments (gold) colored. (E, E’, F, F’) Zoomed-in top views of envelope spikes (orange-red) embedded in the PM with C9 (green) intercalating trimeric spikes. Spikes with bridging C9 density (green) often arranged with clear, near-linear boundaries. Scale bars 25 nm. (G) Subvolume average of budding-arrested NLP below PM and spikes. Docking end of egress-blocked NLPs is flattened in comparison to cytosolic NLP class I (Fig. 3). (H) Plot of distance between C9-linked trimeric spike and nearest neighbor after refinement of orientation for each extracted spike subvolume in representative tomogram shown in (A). Low-resolution subvolume spike average shows general agreement with the CHIKV spike atomic model (PDB:3J0C). (I) Distances on the virus particle between icosahedral-3-fold (i3) spikes and quasi-3-fold spikes (q3), as well as q3-q3 spikes, displayed on the virus particle with icosahedral 2-fold (orange disc), 3-fold (blue triangle) and 5-fold (green penton) for reference.

C9-bound trimeric spikes were readily detected in the tomograms, likely due to condensing of spikes crosslinked by C9, mass addition of bound IgG, and exclusion of other host membrane proteins (Fig. 6E-F, Movie S2). Spike-C9 complexes coalesced into large patches at micrometer scale (Jin et al., 2018). Neither ordered spike assemblies nor direct lateral interactions between spikes, like in spike hexagonal or icosahedral lattices, were observed, while densities between spikes with characteristic Y-shaped features of IgG molecules were readily detected (Fig. 6E-F). The clear boundaries of the coalescence of the spike-C9 complexes, with spikes often lined up, further suggests that spikes are cross-linked and spaced apart via bivalent binding of C9 IgGs. Additionally, subvolume averaging of 7,678 manually-picked, individual spikes yielded a low-resolution density map ∼24 Å (0.143 FSC criterion) that approximately matches the atomic model of the CHIKV trimer (Fig. 6G). Based on the refined spike subvolume orientations, the median distance between nearest-neighbor spikes in the C9-induced coalesced spike patches was determined to be 185.2 Å (range from 98 Å to 309 Å) (Fig. 6H). This distance between centers of C9-linked spikes is mostly greater than that between neighboring spikes in pentagons and hexagons on the surface of mature icosahedral virions (117 Å and 120 Å respectively) (Fig. 6I). This suggests C9 IgGs bridge between spikes and prevent formation of lateral interfaces required to form pentagon and hexagon assemblies on the budding particle surface. The absence of ordered spike assemblies in the spike-C9 coalescence further suggests that spikes are unlikely to be delivered to the PM as pre-assembled lattices, arguing against what was proposed from the observation of hexagonal spike lattice tubes in cytopathic vacuole type-II in Semliki Forest virus-infected cells (Soonsawad et al., 2010).

## Discussion

Here we directly imaged CHIKV-infected cells to gain an enhanced understanding of how Cp and spike proteins coordinate the assembly of icosahedral virus particles *in cellulo* at the molecular level. As illustrated in Figure 7, immature, cytosolic NLPs lacking observable symmetry promote nucleation of an icosahedral spike lattice at the PM that transforms structurally heterogeneous, cytosolic NLPs into expanded, icosahedral viral NC cores during virus budding. Lateral spike interactions progressively form the icosahedral envelope lattice on the surface of budding particles, apparently through sequential addition of individual spikes, rather than from rearrangement of preformed spike arrays. Vertical spike:Cp interactions transmit symmetrical organization from outer- to inner-shell. This outside-to-inside regulation is further supported by the flattened docking end of budding-arrested NLPs apparently re-organized by NAb-crosslinked spikes on the PM surface (Fig. 6B, G). Our model supports both the existing paradigm that spike:Cp and spike:spike interactions are required for alphavirus assembly and a model of alphavirus budding where spike shell assembly provides the driving force for icosahedral particle formation (Fig. 7).

**Figure 7.**
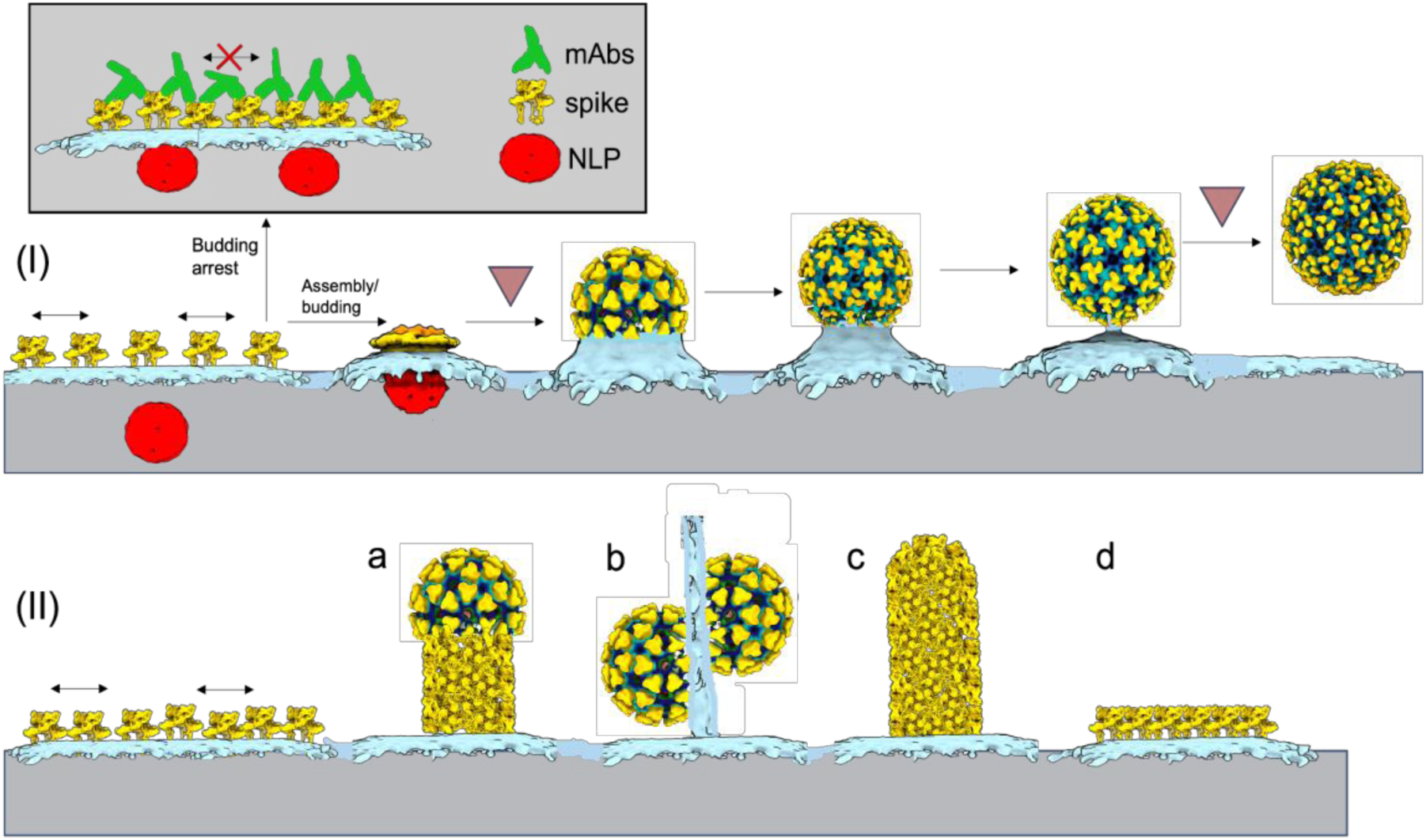
Mechanistic model of alphavirus budding and assembly. At the CHIKV-infected cell surface, immature non-icosahedral NLPs and membrane-embedded spikes converge. Subsequent virus budding (I) is predicated on assembly of the icosahedral spike lattice that enwraps NLPs and reorganizes them into icosahedral NCs through sequential spike:Cp interactions. Rate-limiting steps to particle formation likely occur at early- and late-stages associated with assembly of the first half of the virions and membrane scission following completion of full virions, respectively (upside-down red triangles). Released virions contain near-icosahedral spike and NC layers, with local disruptions in the lattices likely related to membrane scission and virus release from the PM. Binding of mAbs to exposed spike surfaces at the PM (boxed) inhibits virus biogenesis by preventing formation of the curved, icosahedral spike shell. (II) Spikes can self-assemble into non-icosahedral structures, giving rise to alternative assembly products, including (a,c) helical tubes formed by spike hexagons, (b) thin extensions of linked, incomplete particles and (d) planar hexagonal sheets of spikes.

In the CHIKV-infected cell system, we were able to capture snapshots of virus budding from the PM, showing many distinct assembly states. While cryoEM is now commonly used to produce 3D reconstructions, the single-molecule nature of imaging also allows for analysis of a macromolecule’s entire conformational landscape at atomic detail through static images of its many individual states. Analysis of CHIKV assembly states through the entire progression of virion budding differs significantly from past *in situ* cryoET and subvolume averaging studies of non-enveloped viruses with apparently distinct assembly intermediate populations in the cells (Dai et al., 2013; Sutton et al., 2020; Vijayakrishnan et al., 2020). Our discrete-state method of classifying and averaging those individual budding states with compositional and conformational heterogeneity into ensemble 3D classes is an important step in studying progression of a transient assembly process on the cell membrane. In addition to capturing assembly intermediates which are not amenable to *in vitro* purification, our study shows the usefulness of using cryoET to directly derive models of dynamic biological processes in the cell. From the population of particles in each of the 12 budding classes (Fig. 2), it was possible to derive a model of non-uniform budding progression. This model suggests that formation of the first half of the budding shell around the NC, and final scission of the budding membrane neck to release a fully-assembled virion, are likely energetically unfavorable processes and rate-limiting steps in the budding process *in situ*.

In analyzing the contribution of spikes and NCs to assembly of icosahedral particles, our study suggests cytosolic NLPs are roughly spherical but lack pre-formed symmetric organization and therefore do not directly guide icosahedral placement of spikes at the PM during virus budding. Instead, NLPs promote efficient budding of infectious particles by serving as a rough scaffold with suitable curvature to guide assembly of the icosahedral spike lattice. This function in controlling particle size and architecture is reminiscent of dsDNA bacteriophage scaffolding proteins that control the hexamer:pentamer ratio during icosahedral capsid assembly (Chen et al., 2011). Our discovery conflicts with the Cp-centric alphavirus assembly/budding model derived from studies with biochemically reconstructed CLPs (Cheng and Mukhopadhyay, 2011; Ferreira et al., 2003; Mukhopadhyay et al., 2002; Snyder et al., 2011; Wang et al., 2015). Future studies examining early assembly of Cp-gRNAs into cytosolic NLPs and intracellular NLP trafficking using focused ion beam milling (Rigort et al., 2012) of infected cells can provide additional insights into NC morphogenesis. Interestingly, asymmetric NCs serving as a scaffold for spike-driven virus budding can be a common mechanism of particle formation among flaviviruses and alphaviruses (Ferlenghi et al., 2001; Tan et al., 2020).

Our study for the first time demonstrated that self-assembly of spikes produced rare, alternative hexagonal lattices on the PM *in situ* (Fig. 4B-C), in addition to driving budding of the predominant icosahedral virions (Fig. 4A). The hexagonal spike lattice coated vesicles shed off from virus-infected cells (Fig. S7D), may function like the capsidless subviral particles formed by transmembrane glycoproteins of other virus families (Allison et al., 2003; Bruss and Ganem, 1991; Ferlenghi et al., 2001; de Haan et al., 1998; Heilingloh and Krawczyk, 2017; Stange et al., 2008; Wang et al., 2009). In an infected host, subviral particles may help infectious virus particles escape from host immune responses by absorbing specific antiviral antibodies as virus decoys (Heilingloh and Krawczyk, 2017; Vaillant, 2021). The long extensions from the cell periphery, in form of helical spike tubes or thin strings of incomplete particles might promote virus cell-to-cell transmission like reported for retroviruses (Nikolic et al., 2011; Sherer and Mothes, 2008; Sowinski et al., 2008), although using different mechanisms. Infection from the incomplete particle strings, or often-observed released multi-core particles, would result in high local multiplicity of infection (MOI) that usually promotes virus infection by saturating host restriction factors (Bieniasz, 2004; Yan and Chen, 2012), and contributes to modulation of viral pathogenesis by maintaining genetic diversity (Sanjuán, 2021; Vignuzzi and López, 2019). Future functional studies of alternative assemblies of alphaviruses, both *in vitro* and *in vivo*, are warranted.

We previously reported that anti-alphavirus NAbs are able to inhibit virus release in addition to their classical function in neutralizing virus entry, and this anti-release function depends on bivalent binding of NAb IgGs to viral spikes (Fox et al., 2015; Jin et al., 2015; Williamson et al., 2021). Spikes crosslinked by NAbs coalesce into membrane patches that prevent membrane envelopment around attached NLPs, therefore blocking virus budding (Jin et al., 2018). Based on these previous studies we proposed that alphavirus assembly/budding is a potential new target for antiviral therapies. Detailed molecular mechanisms of spatially- and temporally-regulated assembly of alphavirus particles in the cellular environment, as well as the mechanism of NAb-induced alphavirus budding inhibition, will facilitate the design of novel anti-budding therapies. In the current study, we for the first time characterized *in-situ* orchestrated assembly of two-layer icosahedral particles driven by lateral spike interactions and demonstrated that intercalation of NAbs between spikes prevented lateral spike-spike interactions required to assemble hexagonal or icosahedral lattices (Fig. 6). Although we discovered the anti-budding functions for anti-CHIKV antibodies using NAbs binding to E2 (Fox et al., 2015; Jin et al., 2015), the detailed mechanism revealed in this study suggests that any intercalating molecule that locks spikes in a conformation preventing lateral spike-spike interactions could inhibit alphavirus budding. Targeting conserved regions exposed on individual spikes with antibody or other cross-linking molecules can serve as pan-alphavirus antivirals without the need to neutralize virus entry. Two recent studies published while this work is in preparation support our hypothesis (Kim et al., 2021; Williamson et al., 2021), where cross-reactive non-neutralizing mAbs targeting the conserved region in alphavirus E1 are able to inhibit virus budding from arthritogenic to encephalitogenic alphaviruses and provide *in vivo* pan-protection against alphavirus infection.

It is also conceivable that blocking the spike lateral interfaces that are exposed on the individual spike surface, but concealed in spike lattices, can also prevent icosahedral spike lattice formation and subsequent virus budding. In addition to IgG-like large biomolecules, small molecules designed to accurately target spike-spike interfaces can potentially be developed as anti-alphavirus therapies. In the current study we demonstrated that spike-Cp vertical interactions initiate the icosahedral virus assembly/budding. Interestingly, microinjection of synthesized peptide corresponding to the E2 cytoplasmic domain that interacts with NC was reported to successfully inhibit SINV and SFV budding (Kail et al., 1991). It is conceivable that membrane penetrating molecules that interfere with spike-Cp interactions could functionally inhibit alphavirus biogenesis and spreading. Targeting both the extracellular domain and intracellular tail of spikes concurrently will reduce the chance of escaping for alphaviruses that have high mutation rate and often escape from antivirals quickly. Taken together, our current study provides valuable structural information on developing intervening molecules that block alphavirus assembly/budding in two ways: preventing lateral interactions between spikes from outside and/or uncoupling NC and spike assembly from inside of virus-infected cells.

## Methods

### KEY RESOURCES TABLE

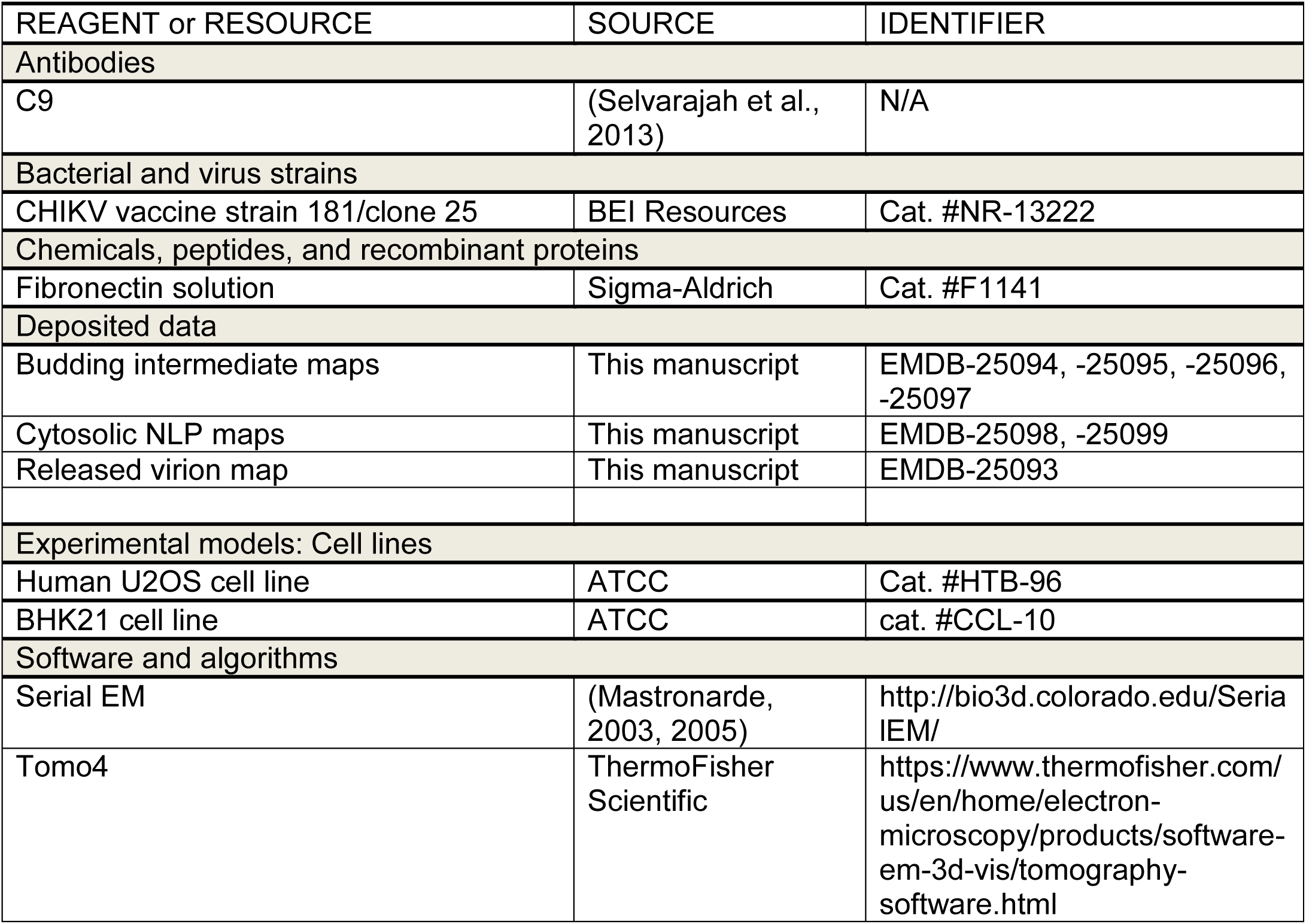
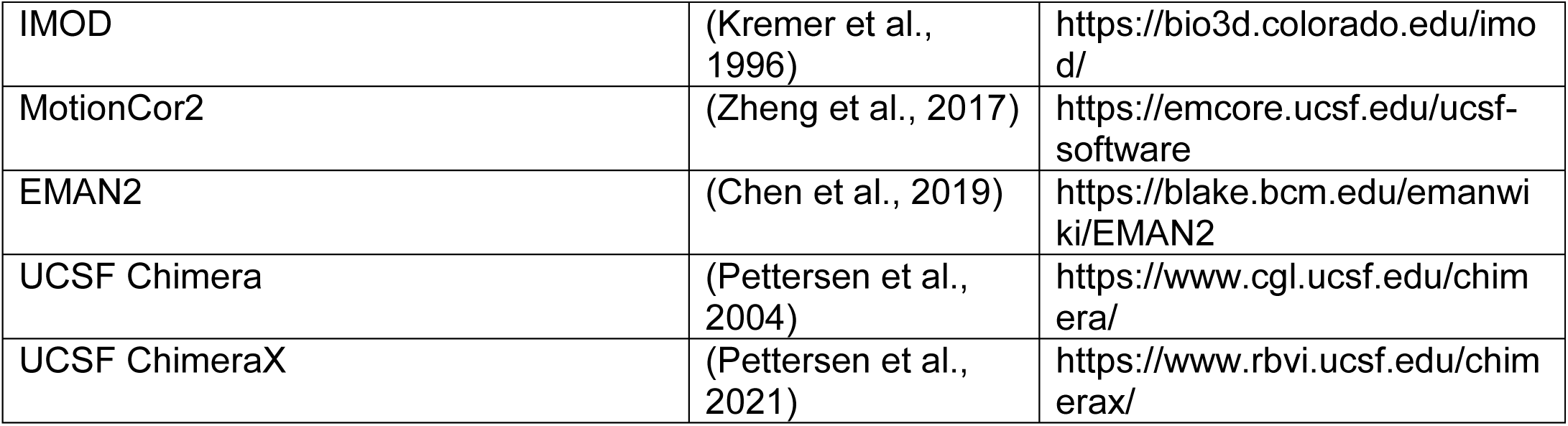

### RESOURCE AVAILABILITY

#### Lead Contact

Further information and requests for resources and reagents should be directed to and will be fulfilled by the lead contact, Wah Chiu, Jing Jin.

#### Data and code availability

Cryo-EM maps reported in this study have been deposited in the Electron Microscopy Data Bank (EMDB) under the following accession codes: EMDB-25093 (released virion), EMDB-25094, -25095, -25096, -25097 (budding intermediates), and EMDB-25098, - 25099 (cytosolic NLPs).

#### EXPERIMENTAL MODEL AND SUBJECT DETAILS

##### Cell cultures

Human bone epithelial cell line U2OS cells (Cat. #HTB-96) is a female cell line purchased from American Type Culture Collection (ATCC). Hamster fibroblast cell line BHK21 cells (Cat. #CCL-10) were purchased from ATCC. Cells were maintained at 37°C with 5% humidified CO_2_ in DMEM (Invitrogen) supplemented with penicillin and streptomycin, 10 mM HEPES, non-essential amino acids, and 10% FBS (Hyclone). CHIKV vaccine strain 181/clone 25 (CHIKV-181) was amplified in BHK21 cells.

##### Virus Strain

CHIKV vaccine strain 181/clone 25 (CHIKV-181) was amplified in BHK21 cells.

## METHODS DETAILS

### Cell infection and vitrification

U2OS cells grown on fibronectin-coated gold 200 mesh R2/2 grids (Quantifoil) were infected with CHIKV-181 at an MOI of 50 for an incubation period of 8 hrs. In the case of neutralizing antibody-treated cells, after 3 hrs of infection, the grids were washed extensively and incubated with 5 ug/mL NAb C9 for an additional 5 hrs. The grids were then washed with PBS and a solution of 10 nm BSA gold tracer (Cat. #25486, EMS) was added directly prior to vitrification. Grids were blotted and plunged into liquid ethane using the LEICA EMGP plunge freezer device. Grids were stored under liquid nitrogen conditions until required for data collection.

### Acquisition and processing of cryo-ET tilt series

Grids of vitrified virus-infected cells were imaged on two instruments: (1) a Titan Krios microscope (ThermoFisher) operated at 300kV with post-column energy filter (20eV) and K2 Summit detector (Gatan) with a calibrated pixel size of 2.72Å and (2) a Talos Arctica (ThermoFisher) operated at 200kV with post-column energy filter (20eV) andK2 Summit detector with calibrated pixel size of 3.54Å. Single-axis, bi-directional tilt series were collected using SerialEM software with low-dose settings and defocus range of -3 to -5.5 µm. For data of CHIKV-181-infected cells collected with the Titan Krios, a total cumulative dose of 110e^-^/A^2^ was applied to the specimen, while for data collected with Talos Arctica, the total average dose at the specimen was 90e^-^/A^2^. In both cases the electron dose was distributed over 51 tilt images, covering an angular range of -50° to +50**°**, with an angular increment of 2°. Additional data collection on both electron microscopes was collected using a Volta phase plate, whereby the objective aperture was removed, phase plate inserted and activated, and tilt series collected under the above conditions. The activated Volta phase plate was operated at phase shift 0.3-0.6π radians as measured by AutoCTF software (ThermoFisher). The motion between frames of each tilt image in the tilt series was corrected using MotionCor2 software (Zheng et al., 2017). Tilt images were compiled, automatically aligned and reconstructed using EMAN2 software (Chen et al., 2019). In total, 144 tomograms were judged as sufficient for further analysis from the Titan Krios data collections and 20 tomograms from the Talos Arctica data collections. A summary of the Cryo-ET data collection can be found in Supplementary Table 1.

For analysis of CHIKV-181-infected cells treated with NAb C9, 61 single-axis, bi-directional tilt series were collected on the Titan Krios microscope operated at 300kV with post-column energy filter and K2 Summit detector and calibrated pixel size of 2.72Å. Data was acquired using SerialEM software with low-dose settings and defocus range of -3 to -5.5 µm. Tilt series were collected with a total cumulative electron dose of 120e^-^/A^2^ distributed over 51 tilt images, again covering an angular range of -50° to +50**°**, with an angular increment of 2°. Data was exclusively collected using an activated Volta phase plate, with phase shift targeted in the range 0.3-0.6π radians. 51 tomograms were judged as sufficient for further analysis, based on achieved phase shift and tomogram reconstruction quality, and were used for subvolume analysis.

### Classification of budding intermediate subvolumes

Subvolume analysis steps were performed using the EMAN2 Tomo pipeline (Chen et al., 2019). CTF estimation for each tilt image was performed using the EMAN2 program *e2spt_tomoctf*.*py*. 1,918 budding intermediate particles were manually picked using the EMAN2 3D slice picker and extracted into subvolumes with x4, x2, x1 binning. 50 high-SNR particles (x4 binning) were picked from the dataset for each of three rough stages of budding (early-, mid-, and late-) for initial model generation. The initial model for each budding class was produced using the EMAN2 initial model generation program *e2spt_sgd*.*py*, first imposing c1 symmetry and running 5 iterations. After aligning the C1 initial models to the symmetry axes, 5 additional iterations were run with C5 symmetry imposed for each. These three maps were then used as initial models for subtomogram multi-reference refinement (*e2spt_refinemulti*.*py*).

The full dataset of 1,918 budding-intermediate subvolumes (x4 binning) was input into EMAN2 multi-reference refinement with 10 initial models (three copies of early-, three copies of mid-, and four copies of late-budding models described above) and run for 12 iterations, imposing C5 symmetry and limiting resolution to 40Å for alignments. Due to poor convergence of the earliest-budding classes, all budding particles in the tomograms were re-picked with two points defining an initial budding orientation: one at the center of NC and one at the apex of the budding shell. Multi-reference refinement of the pre-oriented subvolumes was repeated as described above, with a refinement angular difference constraint of 30° to prevent particle “flipping” from the initial and rough budding orientation. If a resulting class displayed budding virus structural features with sufficient particle count, those particles were subjected to further classification with either two or three low-passed versions of the class average as initial references. In this way, particles within five of the 10 3D classes were subjected to a second round of multi-reference refinement for further identification of budding conformations, with refinement parameters described above (Fig. S3). Between the two rounds of classification, 12 different 3D budding structures were determined in total. Subvolume particles within “junk” class averages lacking interpretable structure were viewed in the original 3D tomograms, revealing these particles covered a wide range of budding levels and were typically located near high density gold fiducials that biased the alignment.

### Subtomogram averaging of budding intermediates, released virions and NLPs

For each of the five budding intermediate 3D classes displaying low-resolution icosahedral features, particles were re-extracted (x4, x2, x1 binning) for subtomogram refinement (*e2spt_refine*.*py*). For each class, 4-6 iterations of refinement were performed for each binned (x4, x2, x1) particle set, imposing C5 symmetry at each step and following gold-standard protocol: all particles were split into two independent subsets and resolution measured by Fourier shell correlation (0.143 FSC criterion) of the two density maps. Following subtomogram refinement of the least-binned particle set for each class, 2 iterations of sub-tilt refinement (*e2spt_tiltrefine*.*py*) with imposed C5 symmetry were performed to produce final budding-intermediate subvolume averages. A summary of the CryoET data collection and subtomogram analysis of viral intermediates can be found in Table S1.

Subtomogram averaging of released virions was performed by manually picking and extracting 521 released particles (x4, x2, x1 binning) into subvolumes, followed by EMAN2 3D refinement and sub-tilt refinement. An initial reference for 3D refinement was generated from 50 high SNR particles with different defocuses using EMAN2, with C5 symmetry imposed as described previously. 3D refinement was performed with C5 symmetry imposed, working from x4 to x2 to x1 binned subvolumes as resolution improved. After visual observation of icosahedral structure in the map, icos. symmetry was applied during final sub-tilt refinement of x1 binned subvolumes. This resulted in a converged map with pixel size 2.72 Å/pixel and resolution (0.143 FSC criterion) of 8.2 Å.

For subtomogram averaging of NLPs, 545 NLPs apparently within the cytosol of virus-infected cell tomograms were manually picked using the EMAN2 3D slice picker and extracted (x4binning) into subvolumes. 50 high SNR particles with varying defocus were used to generate an initial reference with C5 symmetry as described above. Multi-reference refinement of the 545 NLPs (x4 binning) was performed with three classes and similar refinement parameters described above for budding intermediate classification, without the angular difference constraints. This resulted in two cytosolic NLP 3D classes (class I & II) with interpretable structure (Fig. S3). Additional 3D refinements of particles within those two respective classes, with imposed C5 symmetry, resulted in maps with resolutions of 47.6Å (class I) and 43.5Å (class II) (Gold-standard, 0.143 FSC criterion). The refined orientations of cytosolic NLPs within one class displaying local five-fold symmetry (class II, Fig. 3, S4) were mapped back in 3D to originating tomogram reconstructions using EMAN2 program *e2spt_mapptclstotomo*.*py*.

### Subvolume analysis of NAb-crosslinked spikes and budding-arrested NLPs

For analysis of the C9-treated CHIKV-181-infected cells, 7,678 individual spikes were automatically picked from tomograms based on a low-resolution reference and judged individually for false positives. Any additional spikes in the tomogram were picked manually. This extensive manual picking protocol was meant to ensure all spikes were properly extracted for nearest-neighbor distance analysis. 3D subvolumes (x4, x2 binning) of each spike were then extracted and a c3-symmetric initial model was built from a subset of 500 (x4 binning) high SNR particles using the reference-free initial model program in EMAN2 (*e2spt_sgd*.*py)*. The full set of 7,678 (x4, x2 binned) spike particles was then subjected to iterative 3D subtomogram refinement (*e2spt_refine*.*py*) with C3 symmetry imposed until no improvement in refined orientations was achieved. The final converged average map had resolution 24.4Å (Gold-standard, 0.143 FSC criterion) and pixel size 5.44Å/pixel. The Euclidean distance between each refined spike and its nearest neighbor in the dataset was determined using the refined center-of-mass orientations of spike subvolumes in each tomogram.

From the same tomograms, 1,727 budding-arrested NLPs were manually picked, extracted into subvolumes (x4 binned) and an initial model was generated from 50 particles in the dataset as described above with C5 symmetry imposed. 3D refinement of the 1,727 NLP subvolumes, imposing C5 symmetry, resulted in a converged map with pixel size 14.16Å/pixel and resolution of 37.1Å (Gold-standard, 0.143 FSC criterion).

Visualization, figure generation and model docking were performed in UCSF ChimeraX and its built-in tools (Pettersen et al., 2021).

## Acknowledgements

We thank SLAC National Accelerator Laboratory for access and support of these studies, and all SLAC cryoEM staff for technical support and assistance. We also thank Dr. Muyuan Chen for helpful discussions and providing technical advice in data analysis. This research was supported by the NIH grants R01AI148382, P01AI120943 and S10OD021600 (to W.C.) and R01AI119056 (to G.S.).

## Author Contributions

D.C., J.J., and W.C. designed the study. D.C and J.J. performed cryoEM sample preparation and collected cryoET data. D.C. performed 3D reconstruction and subtomogram averaging. D.C, J.J., M.S., and W.C. analyzed the data. D.C., J.J., and W.C. wrote the manuscript with support from all co-authors.

## Supplemental Information

**Fig. S1.**
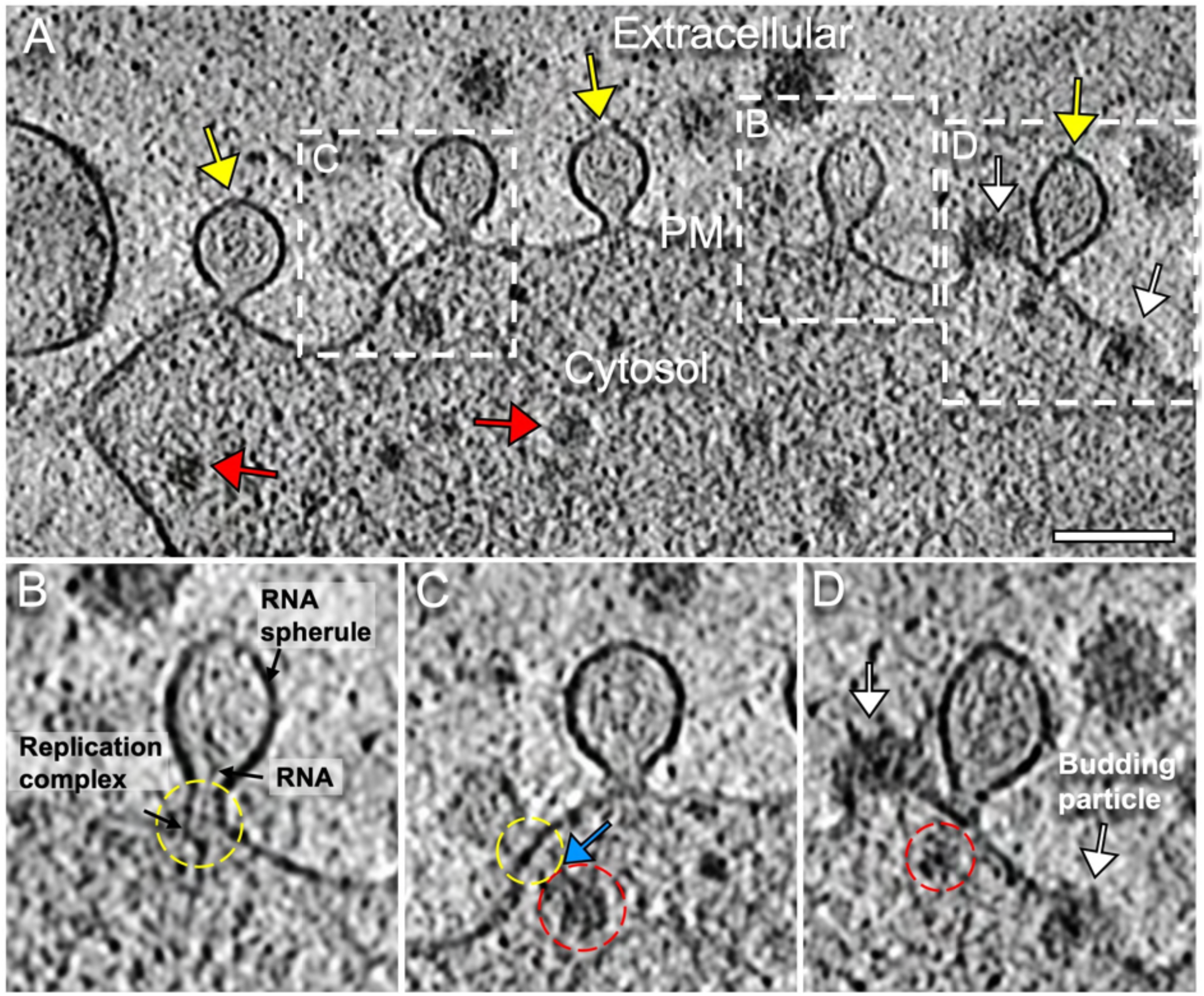
RNA replication spherules on the cell surface. (A) Tomogram slice displaying cell periphery with RNA replication spherules (yellow arrows), budding intermediates (white arrows) and apparently cytosolic NLPs (red arrows). Scale bar: 100 nm. (B) Enlarged tomogram slice of RNA replication spherule with components labeled. Proposed location of the replication complex at the neck of spherules was indicated with a yellow dashed circle. (C) Enlarged slice view of a RNA spherule with a NLP (red dashed circle) in close proximity to the spherule neck (yellow dashed circle), with some thin density (blue arrow) connecting in between. (D) Enlarged slice view of RNA spherule in close proximity to another apparently incompletely-assembled NLP (red dashed circle) and multiple budding particles (white arrows).

**Fig. S2.**
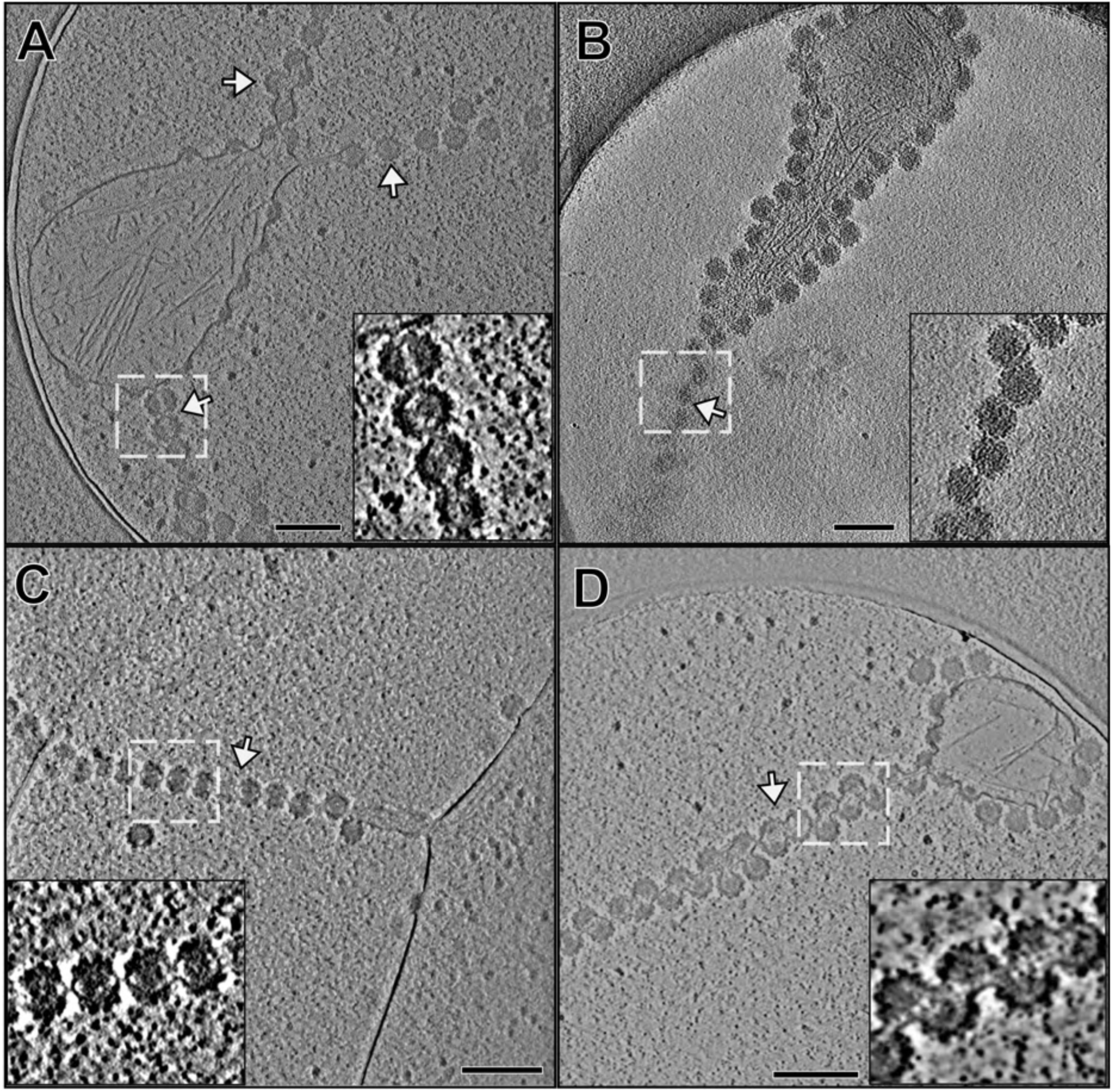
Strands of incomplete particles extend from the cell surface. (A-D) Tomogram slice images display thin extensions of virus budding intermediates in multiple conformations (white dashed boxes, inset images). In all observed cases, beads of linked particles form at the convergences of two membrane surfaces with near-continuous budding intermediates. Scale bars: 200 nm.

**Fig. S3.**
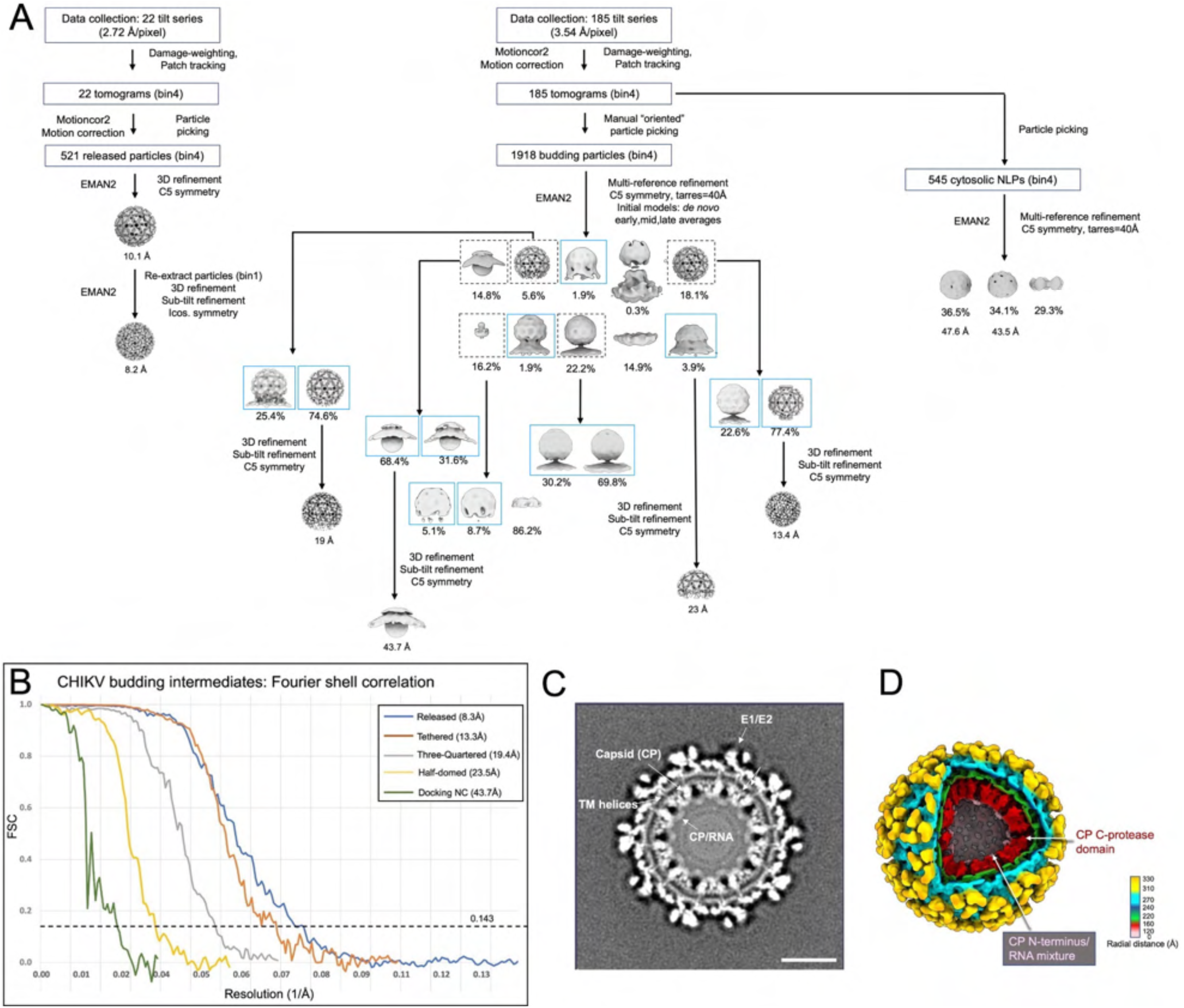
CryoET data processing workflow and resolutions of STA structures. (A) CryoET and data processing workflow. For budding intermediates subvolume classification (middle), particles in 5 of the 10 classes (dashed black boxes) from the first multi-reference refinement were subjected to additional classification. 12 distinct intermediate budding conformations (blue boxes) were resolved after two rounds of classification. Note: two maps displayed within a single blue box merged into one class due to overall structure similarity. (B) Gold standard Fourier shell correlation (FSC) plots of subtomogram average structures of budding intermediates and released virions. (C) Slice view of 3D reconstruction of released CHIKV virions. Subnanometer (8.3Å) resolution of structure is evident by resolved TM helices of E1/E2 in the lipid bilayer. (D) Radial-colored density map of icosahedral CHIKV particle reveals density of CP C-protease domain (red) as well as an ordered density layer below (pink) that is possibly CP N-terminus+gRNA.

**Fig. S4.**
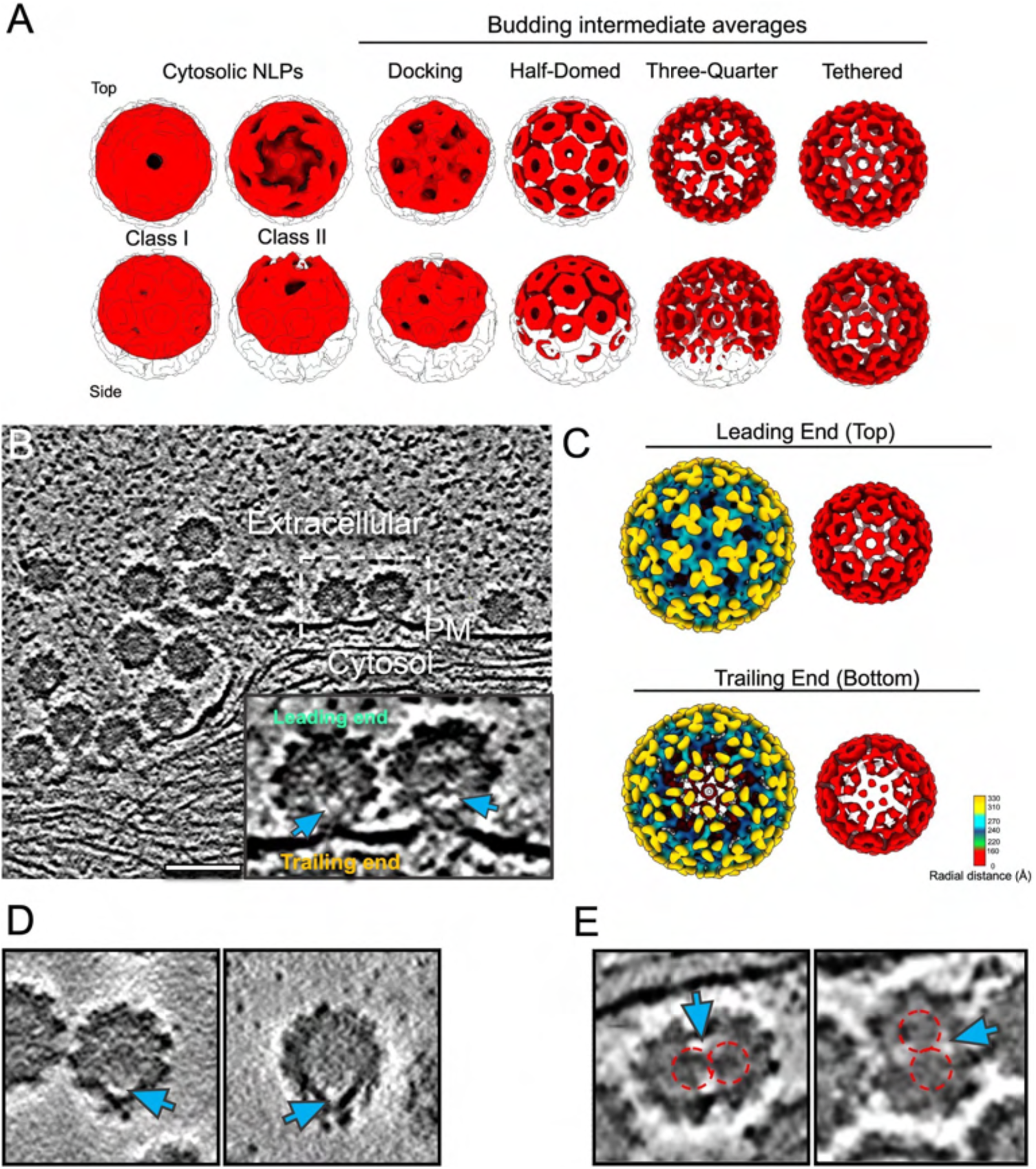
Non-icosahedral features of virus particles. **(A)** Subtomogram average structures of cytosolic NLPs and NCs from budding-intermediate 3D reconstructions (shown in red). NC structures are overlaid on the density map of the NC from released CHIKV virion (white, transparent). (**B**) Volta phase plate tomogram slice images of PM with multiple budding intermediates, including late-stage (“tethered”) particles. Inset is the zoom-in view of the boxed area in (B) displaying relatively absent density at the base of late-stage budding particles (blue arrow). Top of a budding particle, furthest from PM is defined as the leading end, while base of the particle is defined as the trailing end. (**C**) Subtomogram average structure of “tethered intermediate” shows icosahedral symmetry at the leading end, while trailing end of average shows a disordered final penton and distorted capsomer density in those hexamer units below. Relatively absent density between NC and viral envelope (blue arrows) was also observed at (**D**) the non-spherical pole of released virions, and (**E**) the released multi-cored particles. NCs in multi-core particles are labeled with dashed red circles.

**Fig. S5.**
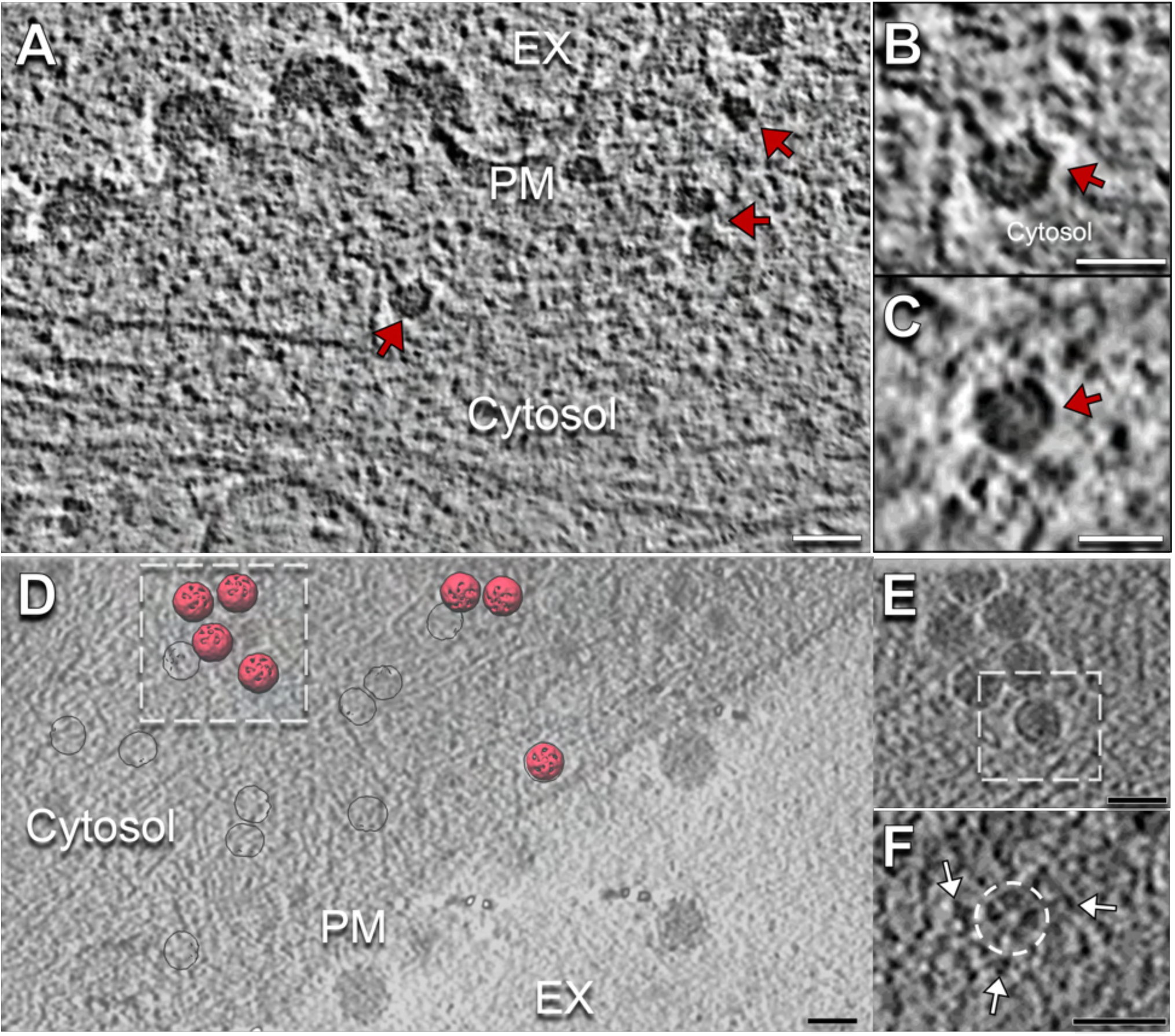
Orientation of cytosolic NLPs. (**A**) Volta-phase plate tomogram slice image shows budding cell periphery with apparently cytosolic NLPs (red arrows). Scale bar: 50 nm. (**B**,**C**) Zoom-in views of NLPs (red arrows). Scale bars: 30 nm. (**D**) Tomogram slice image of cell with subvolume averages of NLP class II (red, Fig. S4) mapped back to the tomogram based on the refined orientation of each particle. 5-fold density consistently oriented towards the PM surface in slices above. (**E**) Tomogram slice image displays cluster of NLPs (red arrow). Scale bar: 50 nm.(**F**) In a tomogram slice directly above a single NLP (white box), a penton of spikes (dashed white circle) is identified along with nearby spikes (white arrows). Scale bar: 50nm.

**Fig. S6.**
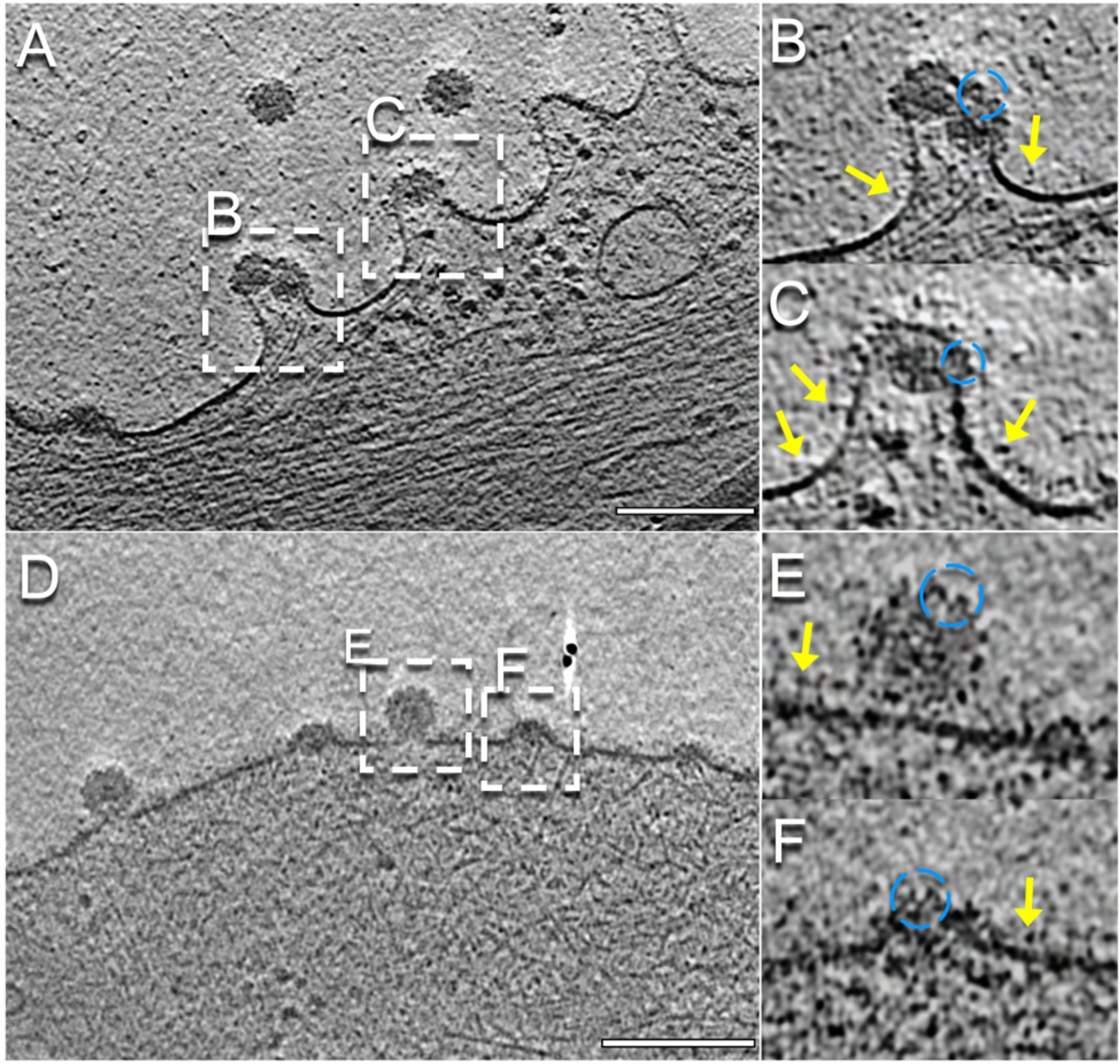
Different conformations of spikes before and after assembly in budding shells. (A,D) Tomogram slice displaying cell periphery with budding intermediates. Scale bars: 200 nm. (B, C, E, F) Enlarged tomogram slices of budding particles. Spike side-views on budding intermediates surface (blue dashed circle) displayed characteristic conformation of spikes assembled in lattice, in contrast to potential individual spikes (yellow arrows) near the base of budding intermediates.

**Fig. S7.**
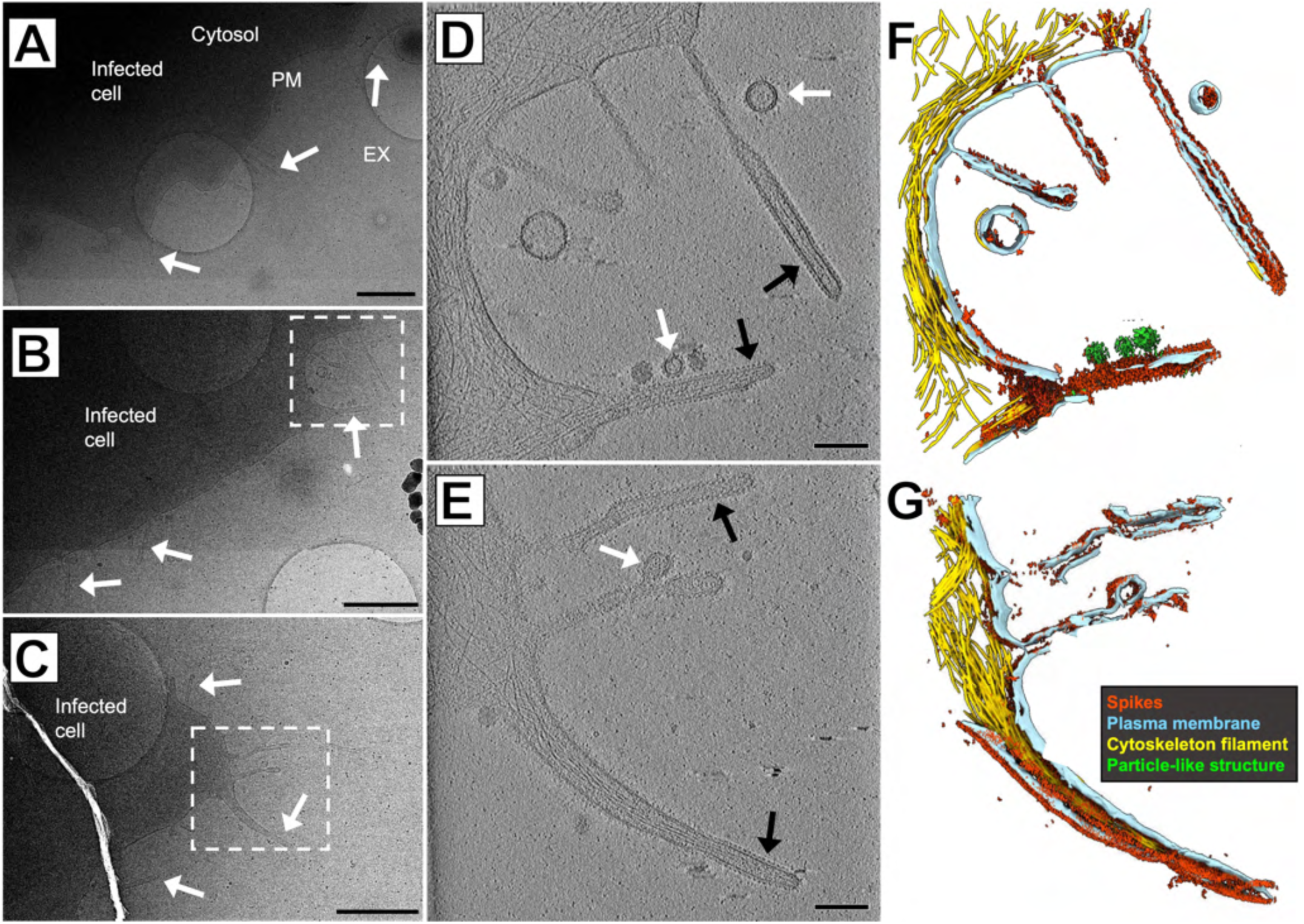
Self-assembled structures of spikes alone. (A-C) Images of CHIKV-infected cell peripheries display cell extensions (white arrows) directly from the cell body. Scale bars: 1 µm. (D & E) Zoom-in images of the boxed regions in B & C, display tubular protein arrays (black arrows) at the terminal end and released vesicle-like assembly products lacking dense cores (white arrows). Scale bars: 200 nm. (F & G) 3D segmentations of cellular features corresponding to (D & E) show the surfaces of tubular extensions contain dense spikes. Segmentations reveal spikes on the surface of tubular membrane extensions that largely exclude bundled cytoskeleton filaments.

**Table S1.**
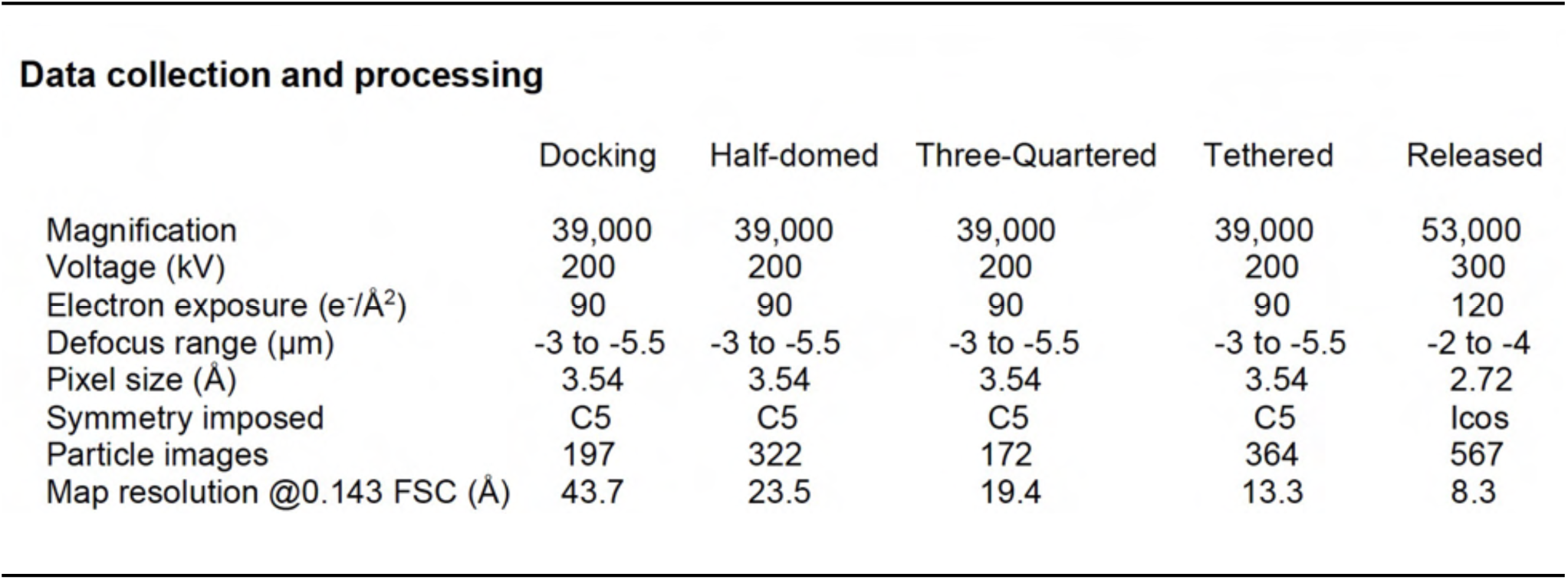
Cryo-ET data analysis and subtomogram averaging statistics.

**Data collection and processing**
|  | Docking | Half-domed | Three-Quartered | Tethered | Released |
| --- | --- | --- | --- | --- | --- |
| Magnification | 39,000 | 39,000 | 39,000 | 39,000 | 53,000 |
| Voltage (kV) | 200 | 200 | 200 | 200 | 300 |
| Electron exposure (e<sup>-</sup>/Å<sup>2</sup>) | 90 | 90 | 90 | 90 | 120 |
| Defocus range (μm) | -3 to -5.5 | -3 to -5.5 | -3 to -5.5 | -3 to -5.5 | -2 to -4 |
| Pixel size (Å) | 3.54 | 3.54 | 3.54 | 3.54 | 2.72 |
| Symmetry imposed | C5 | C5 | C5 | C5 | lcos |
| Particle images | 197 | 322 | 172 | 364 | 567 |
| Map resolution @0.143 FSC (Å) | 43.7 | 23.5 | 19.4 | 13.3 | 8.3 |

**Table S2.**
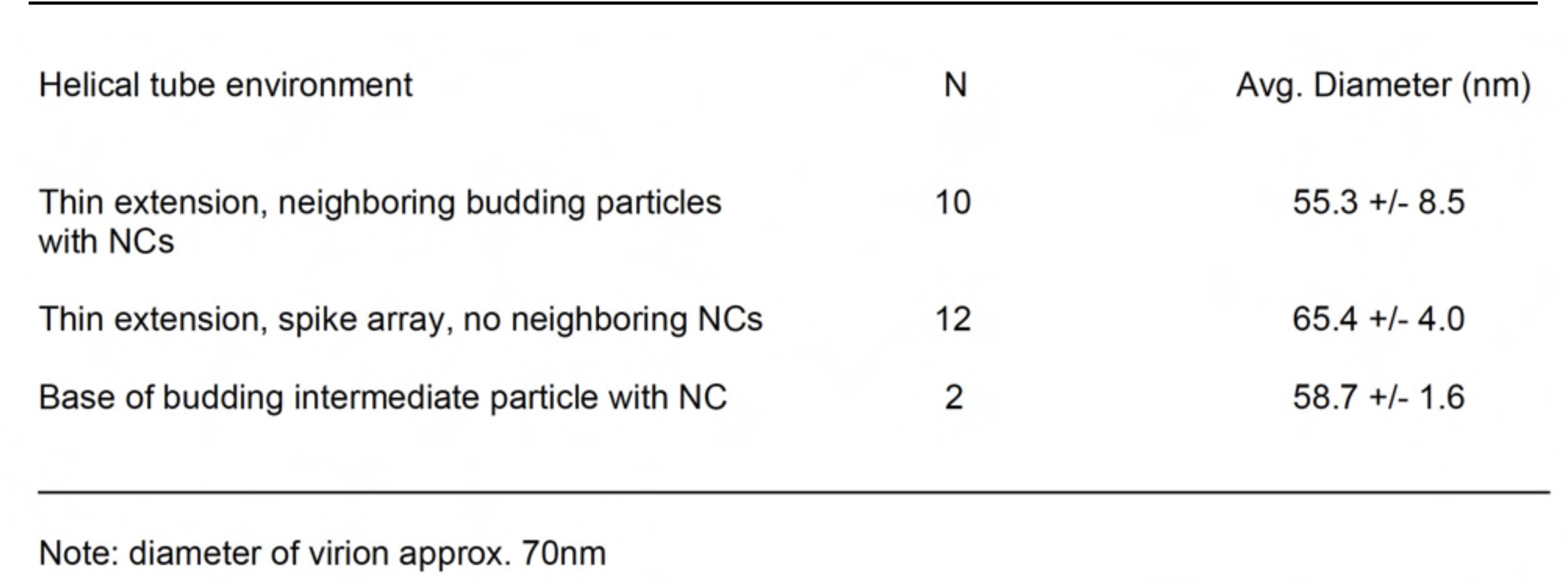
Average diameters of helical tubes formed by hexagonal spike arrays in different cellular contexts.

| Helical tube environment | N | Avg. Diameter (nm) |
| --- | --- | --- |
| Thin extension, neighboring budding particles with NCs | 10 | 55.3 +/- 8.5 |
| Thin extension, spike array, no neighboring NCs | 12 | 65.4 +/- 4.0 |
| Base of budding intermediate particle with NC | 2 | 58.7 +/- 1.6 |

Note: diameter of virion approx. 70nm

